# The innate cytokine IL-18 inhibits CNS autoimmunity through preferential activation of protective CD8 T-cells

**DOI:** 10.64898/2026.01.26.701759

**Authors:** Jeremy A. Morrissette, Vinh Dang, Leonardo Huang, Jemy Varghese, Zachary R. Lanzar, Anastasia Frank-Kamenetskii, Scott W. Canna

## Abstract

Chronic innate immune activation is widely thought to challenge self-tolerance. IL-18 is an inflammasome-activated cytokine and potent amplifier of T-cell activation whose excess is associated with certain autoinflammatory, but not autoimmune, diseases. We tested how excess IL-18 affected susceptibility to experimental autoimmune encephalomyelitis (EAE), a model of CNS autoimmunity driven by IL-18 responsive, myelin-autoreactive CD4 T-cells (CD4T_auto_). We hypothesized that IL-18 would exacerbate immunopathology in EAE. Instead, excess IL-18 was profoundly protective. IL-18 did not impair CD4T_auto_ priming or early expansion. Rather, it selectively restricted later accumulation of highly activated, splenic CD4T_auto_ bearing CNS-homing integrins. Despite high IL-18 receptor expression on CD4T_auto_ and Foxp3^+^ CD4T_reg_, excess IL-18 acted specifically through mature CD8 T-cells to promote a highly activated CD8T_effector_ phenotype and IFNγ-dependent protection from EAE. Therapeutic administration of a “decoy-resistant” IL-18 agonist (DR-18) to wild-type mice, even after CD4T_auto_ expansion, nevertheless engaged CD8 T-cells to diminish CD4T_auto_ abundance, prevent CNS infiltration, and block immunopathology. Together, these findings demonstrate IL-18’s unexpected, dominant ability to mobilize protective CD8 T-cells against highly activated CD4T_auto_ and protect from CNS autoimmune pathology; illustrating a potential therapeutically relevant mechanism by which autoinflammation actively opposes autoimmunity.

## INTRODUCTION

Autoinflammation refers to primary innate immune activation, whereas autoimmunity refers to pathologic responses to self-antigens. This distinction is relatively recent and was inspired by inflammasome-related disorders where chronic, systemic inflammation is not associated with autoantibodies or other features of autoimmune pathology. Subsequent discoveries suggest these disorders may conspicuously lack overlap with autoimmunity compared with other autoinflammatory diseases (1). Inflammasomes activate and enable the secretion of the cytokines IL-18 and IL-1β. Certain diseases (e.g. NLRC4 gain-of-function, XIAP deficiency) associated with very high IL-18 levels are at risk for the systemic inflammatory state called macrophage activation syndrome (MAS), but not autoimmunity (2). Still’s Disease may straddle the autoinflammation/autoimmunity divide, with IL-18^HI^ individuals following a MAS/autoinflammation-predominant course (3).

IL-18 is produced by certain myeloid and epithelial cells as an inactive cytosolic precursor. Inflammasome-mediated cleavage generates its active form and initiates Gasdermin D pore formation through which IL-18 is released. Most circulating IL-18 is neutralized by an abundant antagonist, IL-18 binding protein (IL-18BP). Unbound “free IL-18” signals through its heterodimeric receptor expressed on NK cells, activated T-cells, and potentially some granulocytes and epithelial cells (2). Functionally, IL-18 is best-known as a lymphocyte “Interferon-γ (IFNg)-inducing factor” and an important contributor to Type 1 host defense in experimental systems (4). However, in specific contexts IL-18 exerts a broader range of effects on T-cells, including promotion of Th2 or Th17 differentiation, supporting skin and gut barriers, modulating inflammation, promoting tissue repair, and enhancing regulatory cell function (5–9).

“Excess IL-18” occurs when levels of mature IL-18 greatly exceed those of IL-18BP. IL-18 overproduction is associated with autoinflammation and MAS, whereas deficiency of IL-18BP in humans is described only in three cases of fulminant hepatitis with unclear MAS overlap (10, 11). Deficiency of IL-18, or inflammasomes in general, does not appear to have a human phenotype in the modern era (12). Mice with excess IL-18 via overproduction (*Il18tg*) or IL-18BP deficiency (*Il18bp^KO^*) show similar responses to stimulation (2, 13–15). In sterile- and virus-induced models of MAS, excess IL-18 amplifies immunopathology through (predominantly) CD8 T-cell activation and subsequent IFNg production (13–18). Supporting the validity of these models, MAS patients also have highly-activated CD38^HI^HLADR^+^ CD8 T-cells and biomarkers of IFNg-activity (19–23). Accordingly, exogenous IL-18 shows promise in potentiating CD8 and Chimeric Antigen-Receptor T cell-mediated tumor clearance (24–26).

The role of excess IL-18 in autoimmunity is unclear. IL-18 levels are modestly elevated in many autoimmune diseases (e.g. rheumatoid arthritis, Sjogren Syndrome, and multiple sclerosis (MS)) (27–32), but appear easily buffered by abundant IL-18BP and may not be pathogenic. Studies of IL-18 in models of autoimmunity focus on deficiency and show modest, context-dependent effects on Th17, Tγδ, and CD4T_reg_ function (5, 33–36). MHC-II is the strongest genetic autoimmunity risk locus (37), implicating CD4 T helper cell autoreactivity as particularly dangerous. The experimental autoimmune encephalomyelitis (EAE) model of MS is a well-characterized system for evaluating competing Th1, Th17, and CD4T_reg_ programs (38). In EAE, immunization against myelin-based peptides drives a Th1/Th17-phenotype in autoreactive CD4 T-cells (CD4T_auto_), which traffic to the central nervous system (CNS), produce cytokines, recruit other inflammatory cells, and ultimately drive demyelination and a progressive, ascending paralysis. The role of IFNg is well-studied but complex, exerting both pro-inflammatory effects and protective functions in EAE (39). Studies of impaired IL-18 production/signaling in EAE showed either no effect or modest protection (40, 41) but the effects of excess IL-18 have not been studied. We sought to understand how excess IL-18 affected the development of autoreactivity and manifestations of autoimmunity, hypothesizing that it would amplify the autoreactive program and exacerbate EAE.

## RESULTS

### Excess IL-18 protects against EAE without signs of hyperinflammation

Following MOG^35-55^ EAE induction, WT mice developed the expected progressive, ascending paralysis after about 14 days but, both *Il18tg* and *Il18bp^KO^* mice were unexpectedly protected from the development and progression of neurologic defects and weight loss **(Figure 1A, B)**. Histological examination of *Il18bp^KO^* spinal cords revealed a marked reduction in the number of inflammatory foci relative to WT mice (**Figure 1C).** Notably, in *Il18bp^KO^* mice, protection was a direct effect of IL-18, as both IL-18 deletion (*Il18^KO^;Il18bp^KO^*) and IL-18R1 blockade abrogated protection from EAE (**Figure S1A-B**). Importantly, neither *ll18tg* nor *ll18bp^KO^* mice showed excessive weight loss, splenomegaly, anemia, thrombocytopenia, or leukopenia indicative of hyperinflammation (**Figure 1A, B, D, E**).

**Figure 1.**
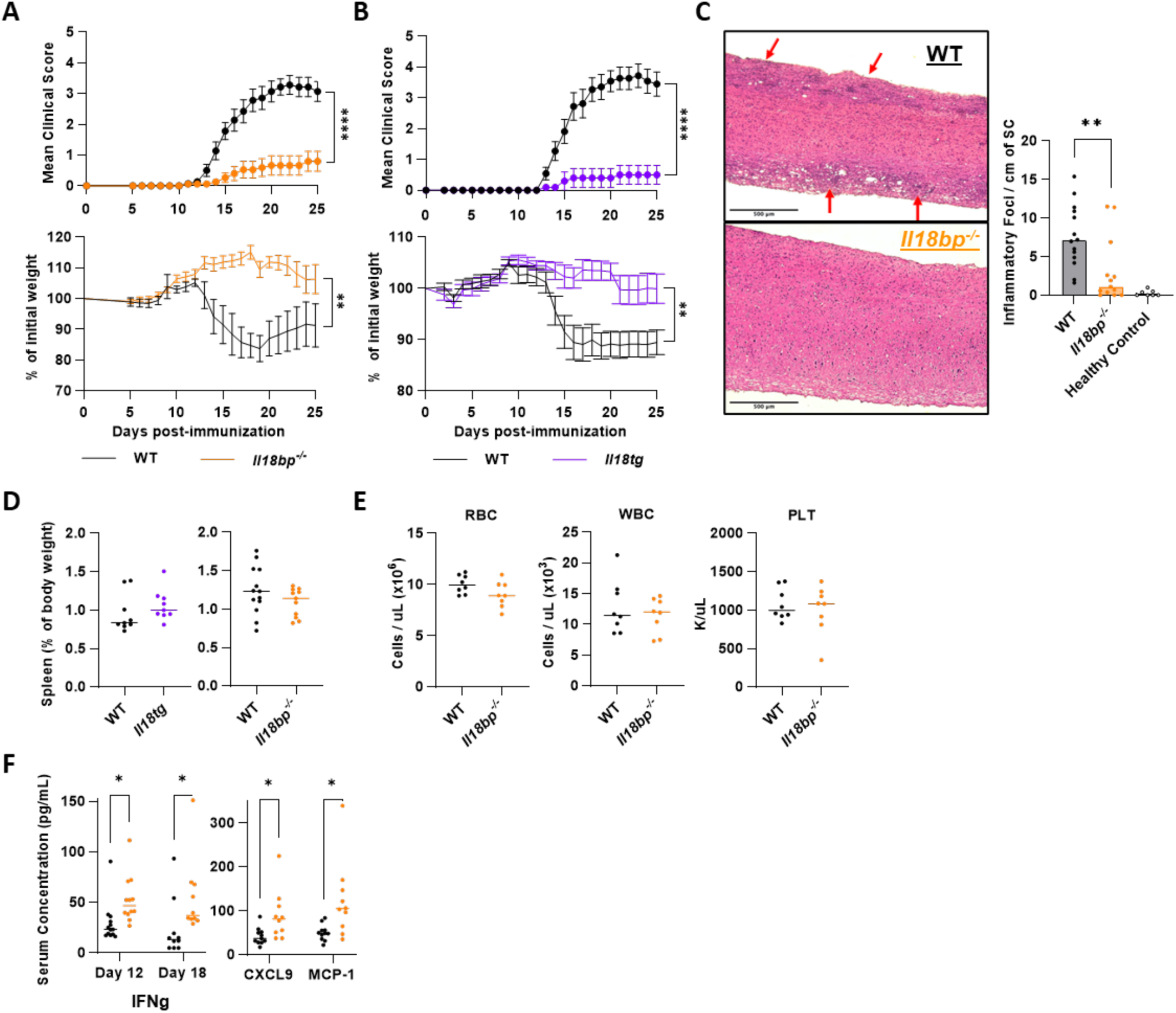
Excess IL-18 protects from EAE without signs of hyperinflammation. **(A)** Mean EAE clinical score and weight change of WT (n=14) vs. *Il18bp^KO^* (n=15) mice or **(B)** WT (n=11) vs *Il18tg* (n=10) mice following MO3^45-55^ EAE induction. **(C)** Representative images and blinded scoring of spinal cord H&E at peak disease (days 18-21). Two sagittal sections per cord were imaged at 50X with inflammatory foci marked by arrows. Number of foci per section were counted and normalized by section length and averaged. **(D)** Spleen weight (normalized to body weight) at days 18-20. **(E)** RBC, WBC, PLT count at day 20. **(F)** IFNg serum levels at day 12 and 18 with IFNg-inducible chemokine serum levels at day 12. (A-F) Data pooled from at least 2 experiments. Error bars = SEM. Statistical analysis: (A,B) Mann-Whitney of area under the curve (AUC) for EAE scores, 2-way RM ANOVA with genotype effect represented for body weight, (C,D) unpaired t-test, (E, F) or unpaired t-tests, p-value with Holm-Sidak correction. Only p<0.05 shown. *p< 0.05, **p<0.01, ****p<0.0001.

Given the canonical role of IL-18 in amplifying IFNg, and the complex involvement of IFNg in EAE pathogenesis, we assessed whether excess IL-18 led to greater levels of IFNg and related cytokines/chemokines during EAE. This was performed in *Il18bp^KO^* mice, as *Il18tg* mice show slightly elevated serum IFNg levels even without stimulation (15, 16). Serum from *Il18bp^KO^* mice showed mildly increased IFNg at day 12 and 18, accompanied by increases in the IFNg-inducible chemokines CXCL9 and MCP-1 at day 12 (**Figure 1F)**. Additionally, homogenate from draining lymph nodes (dLN) of *Il18bp^KO^* mice contained significantly more CXCL9 than WT controls (**Figure S2D)**. These data suggest that, although *Il18bp^KO^* mice are protected from EAE pathology, excess IL-18 induces modestly increased IFNg and downstream cytokines/chemokines.

### Excess IL-18 prevents accumulation of activated CD4 T-cells in the periphery and CNS

We analyzed peripheral immune cells at day 12, roughly the peak of T cell priming and earliest onset of symptoms. *Il18bp^KO^* mice exhibited a modest decrease in splenic cellularity, primarily driven by fewer myeloid cells (CD11b^+^) and CD4 T-cells, including FoxP3+ regulatory T-cells (CD4T_reg_) with no major differences in other populations (**Figure 2A, Figure S2A**). We assessed CD4 T-cell phenotypes and found that *Il18bp^KO^* mice showed a substantial decrease in expression of activation (CD44 and IL-18R1) and inhibitory (PD-1 and CD38) markers (**Figure 2B**).

**Figure. 2.**
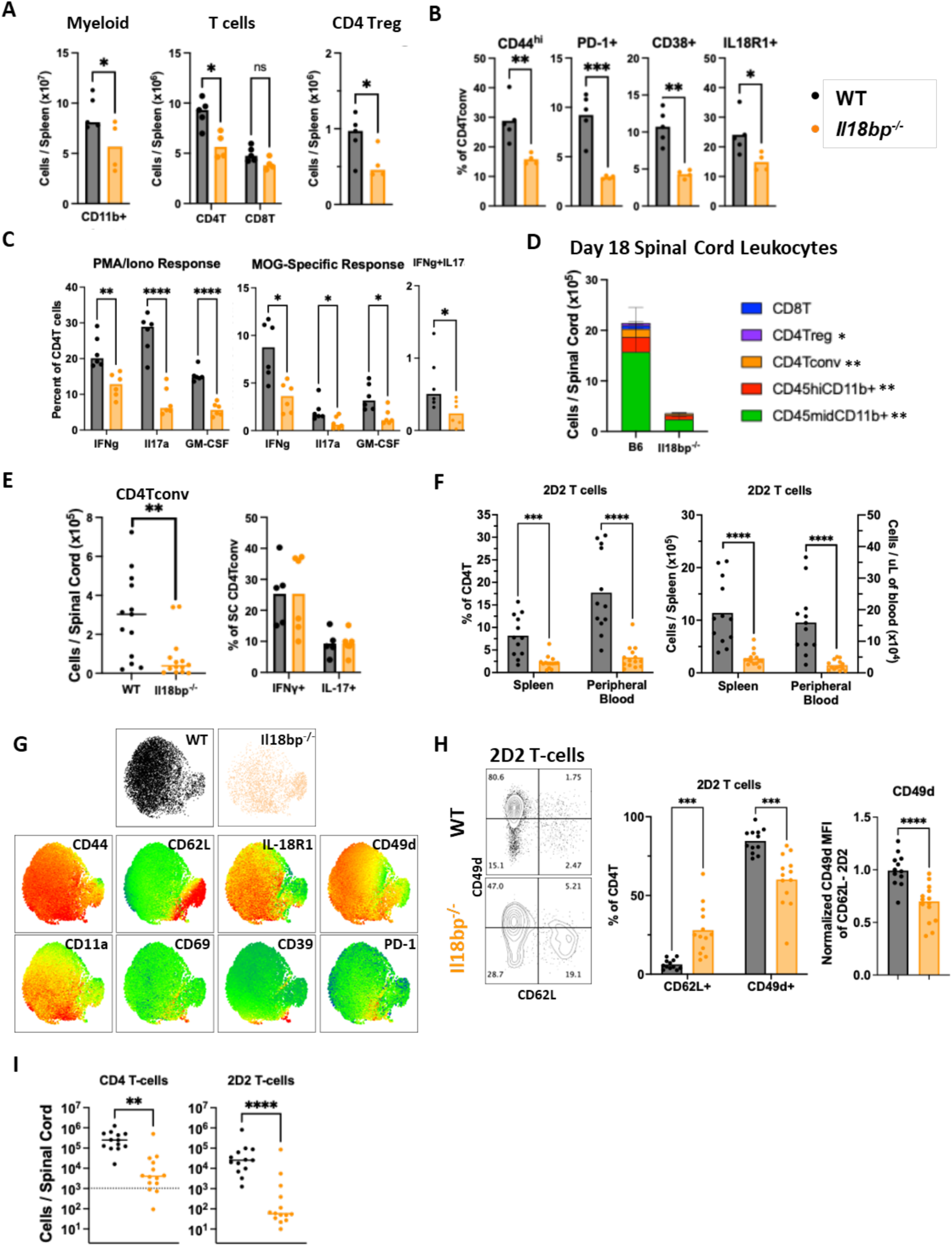
Excess IL-18 prevents accumulation of activated CD4 T-cells in the periphery and CNS. **(A)** Flow cytometric quantification of splenic leukocyte populations in B6 and *Il18bp^KO^* mice at day 12. **(B)** Surface expression of activation markers on splenic CD4Tconv at day 12. **(C)** Quantitation of individual cytokine production by splenic CD4Tconv assessed by intracellular cytokine staining (ICS) following PMA/Iono stimulation and MOG^35-55^ stimulation. Percent of dual-producing IFNg^+^IL-17A^+^ cells following MOG^35-55^ stimulation. **(D)** Flow cytometric quantification of spinal cord leukocytes on day 18. **(E)** Quantification of total spinal cord CD4Tconv and individual cytokine production by ICS after MOG-stimulation at days 18-21. **(F-I)** 2.5×10^5^ CD4 T-cells from naïve 2D2 spleens were transferred to WT and *Il18bp^KO^* mice on day −1 followed by EAE induction on day 0. **(F)** Quantification of 2D2 T-cells at day 12 as a percent and absolute number. **(G)** UMAP visualization of multiparameter flow cytometric data with expression heatmaps of splenic 2D2 T-cells recovered on day 12 from WT and *Il18bp^KO^* mice (see methods for details). **(H)** Representative flow plots and percentage positive of CD62L and CD49d on total splenic 2D2 T-cells at day 12. MFI of CD49d on effector (CD44^+^CD62L^−^) 2D2 T-cells. **(I)** Quantification of spinal cord-infiltrating CD4 T-cells and 2D2 T-cells at day 18. Dashed line represents the average CD4T-cells detected in an unimmunized WT spinal cord. (A, B, C, G) Data representative of 3 experiments. (D, E, F, H, I) Data pooled from 2-3 experiments. Error bars = SEM. Statistical analysis: All data analyzed by unpaired t-tests with Holm-Sidak correction if multiple comparisons made. Only p<0.05 shown. ns = not significant, *p< 0.05, **P<0.01, *** P<0.001, ****p<0.0001.

Secretion of IL-17, IFNg, and GM-CSF have been associated with those autoreactive CD4 T-cells (CD4T_auto_) necessary for driving EAE pathology (42–47). CD4 T-cells from *Il18bp^KO^* mice demonstrated decreased *ex vivo* production of all three cytokines following both non-specific and MOG^35-55^ peptide stimulation. This included the IFNg/IL-17 “double positive” population associated with both EAE and MS (48). Similar trends were seen in peptide-stimulated splenocyte culture supernatants (**Figure 2C, Figures S2B-C**). Th17 cells may undergo microbiota-induced “licensing” in the gut before trafficking to the CNS and driving damage in EAE (42, 49, 50). *Nlrc4^GOF^* mice overproduce IL-18 from intestinal epithelia and show type 1 immune activation in the gut but lack systemic free IL-18 (13, 51). *Nlrc4^GOF^*mice were not protected from EAE (**Figure S2D)**.

Similarly, *Il18bp^KO^* mice had fewer spinal cord leukocytes during periods of CNS involvement (days 18 & 24), including myeloid cells, FOXP3^−^ “conventional” CD4 T-cells (CD4T_conv_), and to a lesser extent CD4T_reg_ (**Figure 2D, Figure S2E-F**). Despite this reduction, those *Il18bp^KO^*CD4T_conv_ identified in the spinal cord maintained IFNg and IL-17 production (**Figure 2E**). Overall, excess IL-18 limited (but did not entirely prevent) the CD4T_auto_ activation and CNS infiltration required for EAE pathology, but it also decreased peripheral and CNS CD4T_reg_.

To specifically assess the impact of excess IL-18 on CD4T_auto_, we performed transfer experiments using “2D2” T-cells derived from a mouse with transgenic expression of a MOG^35-55^-specific TCR (52). 2D2 CD4 T-cells were transferred into WT and *Il18bp^KO^* mice prior to EAE induction. 5 days later, we found equivalent 2D2 proliferation in peripheral lymphoid organs (**Figures S3A-B**), suggesting IL-18 does not alter early antigen-presentation or priming despite its ultimate protective effect on EAE. Transfer of up to 250,000 2D2 T-cells prior to EAE induction did not significantly affect disease in either strain (**Figure S3C**). By day 12, WT mice demonstrated substantial expansion and accumulation of 2D2 T-cells in spleens and blood. By contrast, *Il18bp^KO^*mice failed to accumulate 2D2 T-cells, with nearly five-fold fewer cells in both compartments (**Figure 2F, Figure S3D**).

To better discern differences in transferred 2D2 T-cells, we visualized their surface phenotypes by UMAP (**Figure 2G**). Those recovered from WT mice showed surface markers consistent with highly activated effector status (CD44^hi^, CD62L^−^, IL-18R1^+^) including expression of integrin subunits critical for CD4T_auto_ CNS-infiltration (CD49d and CD11a). By contrast, a smaller proportion recovered from *Il18bp^KO^* mice displayed this same activated effector phenotype, and more cells expressed a “central-memory” phenotype (CD62L^+^, CD49d^−^) **(Figures 2G-H).** Even among effector 2D2 T-cells (CD44^hi^ CD62L^−^), those from *Il18bp^KO^* mice showed lower CD49d expression (**Figure 2H**). Both endogenous and 2D2 CD4 T-cells were reduced in spinal cords of *Il18bp^KO^* mice (**Figure 2I**). The few 2D2 T-cells detected in *Il18bp^KO^* spinal cords showed no differences in per-cell expression of activation markers, including CD49d (**Figure S3E**). Together, these findings suggest that excess IL-18’s effects during the early priming phase of EAE (before day 5) are limited, but at later timepoints excess IL-18 dramatically diminishes the number and activation state of peripheral CD4 T-cells and number of CNS CD4’s.

### IL-18-responsive T-cells mediate protection against EAE

Development and regulation of EAE involves a variety of IL-18 responsive lymphocytes, antigen-presenting, and non-hematopoietic cells (38). To better understand how IL-18 exerts its protective effects, we generated *Il18tg* mice lacking the IL18-receptor specifically on T-cells (*Il18tg;Il18r1*^Δ*T*^). As expected, IL-18R1 was absent on spinal cord and splenic CD4 and CD8 T-cells, but retained on NK cells, from *Il18tg;Il18r1*^Δ*T*^ mice but not *Il18tg;Il18r1^fl/fl^*controls lacking Cre-recombinase expression (**Figure 3A**). T-cell-specific loss of IL-18R completely abrogated the protective effect of excess IL-18 on EAE score (**Figure 3B**), but did not restore the number of splenic CD4T_conv_ or CD4T_reg_ (**Figure 3C**). Intracellular cytokine staining of splenocytes revealed that loss of *Il18r1* in T-cells of *Il18tg* mice restored CD4 T-cell IL-17 production, with minimal effects on IFNg and GM-CSF production (**Figure 3D)**. Spinal cord CD4 T-cells, suppressed in control *Il18tg* mice, were restored to WT levels in *Il18tg;Il18r1*^Δ*T*^ mice (**Figure 3E)**. We observed similar trends in CNS CD4T_reg_, and the opposite trend in CNS CD8 T-cells (**Figure 3E-F**). We did not observe substantial differences in intracellular cytokine staining from CNS CD4 T-cells (**Figure 3G)**. Overall, these data demonstrate that IL-18 acts directly on a population of T-cells to limit EAE and the accumulation of CD4, but not CD8, T-cells in the CNS.

**Figure 3.**
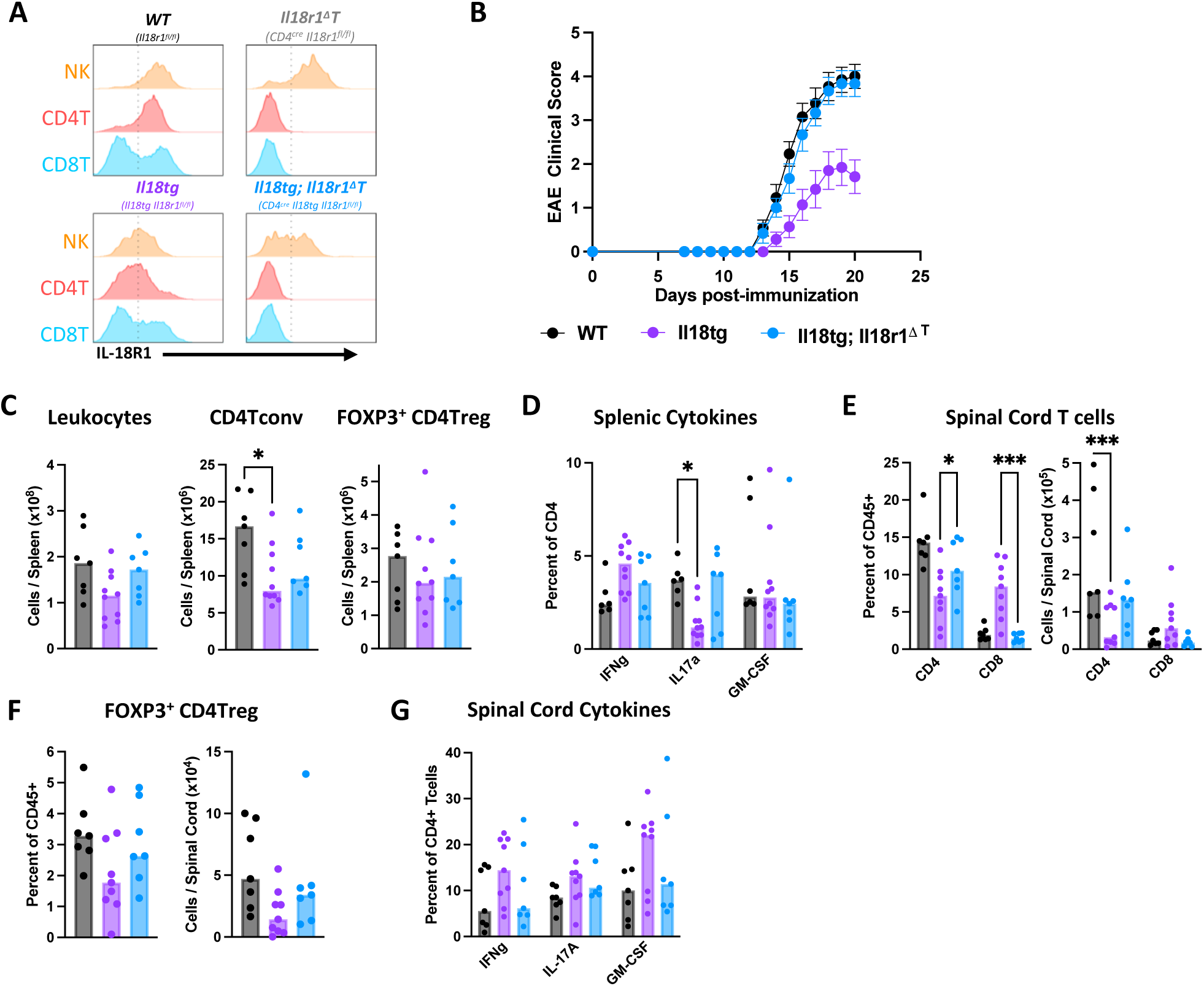
IL-18-responsive T-cells mediate protection against EAE. **(A)** Representative histograms of IL-18R1 expression on NK cells, CD44^+^ CD8 T-cells, and CD44^+^ CD4 T-cells from day 20 dLN. **(B)** Mean EAE clinical score of WT (n=13), *Il18tg* (n=14), and *Il18tg;Il18r1^ΔT^* (n=12) mice. **(C)** Flow cytometric quantification of splenic leukocytes at days 20-21. **(D)** Quantitation of individual cytokine production by splenic CD4 T-cells assessed by ICS following PMA/Iono stimulation. **(E)** Percent and absolute number of T-cells and **(F)** FOXP3^+^ CD4T_reg_ in spinal cords by flow cytometry. **(G)** Quantitation of individual cytokine production by spinal cord CD4 T-cells by ICS following PMA/Iono stimulation. (A) Representative of 3 experiments. Data pooled from (C-G) 2 or (B) 3 experiments. Error bars = SEM. Statistical analysis: (B) Kruskal-Wallis of AUC with Dunns post-test of pairwise comparisons, (C, F) one-way ANOVA with Dunnett’s post-test of pairwise comparisons to *Il18tg*, (D, E, G) 2way ANOVA with Dunnett’s post-test of pairwise comparisons to *Il18tg*. Only p_adj_<0.05 is shown. *p< 0.05, **p<0.01, ***p<0.001.

### IL-18 does not directly inhibit autoreactive CD4 T-cells or protect through FOXP3^+^ CD4T_reg_

We next asked whether IL-18 directly impaired CD4T_auto_, limiting their expansion and CNS-homing, or instead augmented protective FOXP3^+^Tregs during EAE. IL-18 can promote FOXP3^+^ CD4T_reg_-mediated suppression of inflammatory Th17 cells (8), but IFNg is thought to promote CD4T_reg_ fragility (53). To begin testing these possibilities, we generated *2D2;Il18bp^KO^*mice, combining the autoreactive *2D2* TCR with unopposed IL-18. *2D2* mice are highly enriched for CD4T_auto_, but also induce functional FOXP3^+^ CD4T_reg_ during EAE (54). With normal IL-18 regulation, *2D2* mice developed more rapid and severe EAE compared to WT mice, as expected. In contrast to the protection observed in *Il18bp^KO^* mice, *2D2;Il18bp^KO^* instead developed more severe EAE than *2D2* controls (**Figure 4A**). This observation confirmed that IL-18 was not directly cytotoxic to CD4T_auto_, and (at least in *2D2*-transgenic mice) even enhanced pathogenicity. This paradoxical exacerbation of EAE in *2D2;Il18bp^KO^* mice was not attributable to defective CD4T_reg_ differentiation, as all mice showed substantial FOXP3^+^ CD4T_reg_ populations in the periphery and spinal cord (**Figures 4B-C**). Both *2D2* and *2D2*;*Il18bp^KO^* contained very few CD8T-cells during EAE (**Figure 4B**), as expected given allelic exclusion. To further evaluate the relevance of CD4T_reg_ to IL-18’s protection, we examined their phenotypes in WT and *Il18bp^KO^*mice during EAE. A variety of surface markers and transcription factors in CD4T_reg_ showed no significant differences (**Figure S3F-H**). We generated *Il18tg* mice lacking IL-18R1 specifically in FOXP3-expressing CD4T_reg_ (*Foxp3^YFP-Cre^ Il18r1^fl/fl^ Il18tg).* These *Il18r1^ΔFoxp3^*mice did not lose protection from EAE (**Figure 4D**), despite efficient deletion of *Il18r1* specifically in FOXP3^+^ CD4T_reg_ (**Figure 4E, F**). Together, these data demonstrate that excess IL-18 does not exert its protective effect directly through FOXP3^+^ CD4T_reg_ or conventional CD4T_auto_.

**Figure 4.**
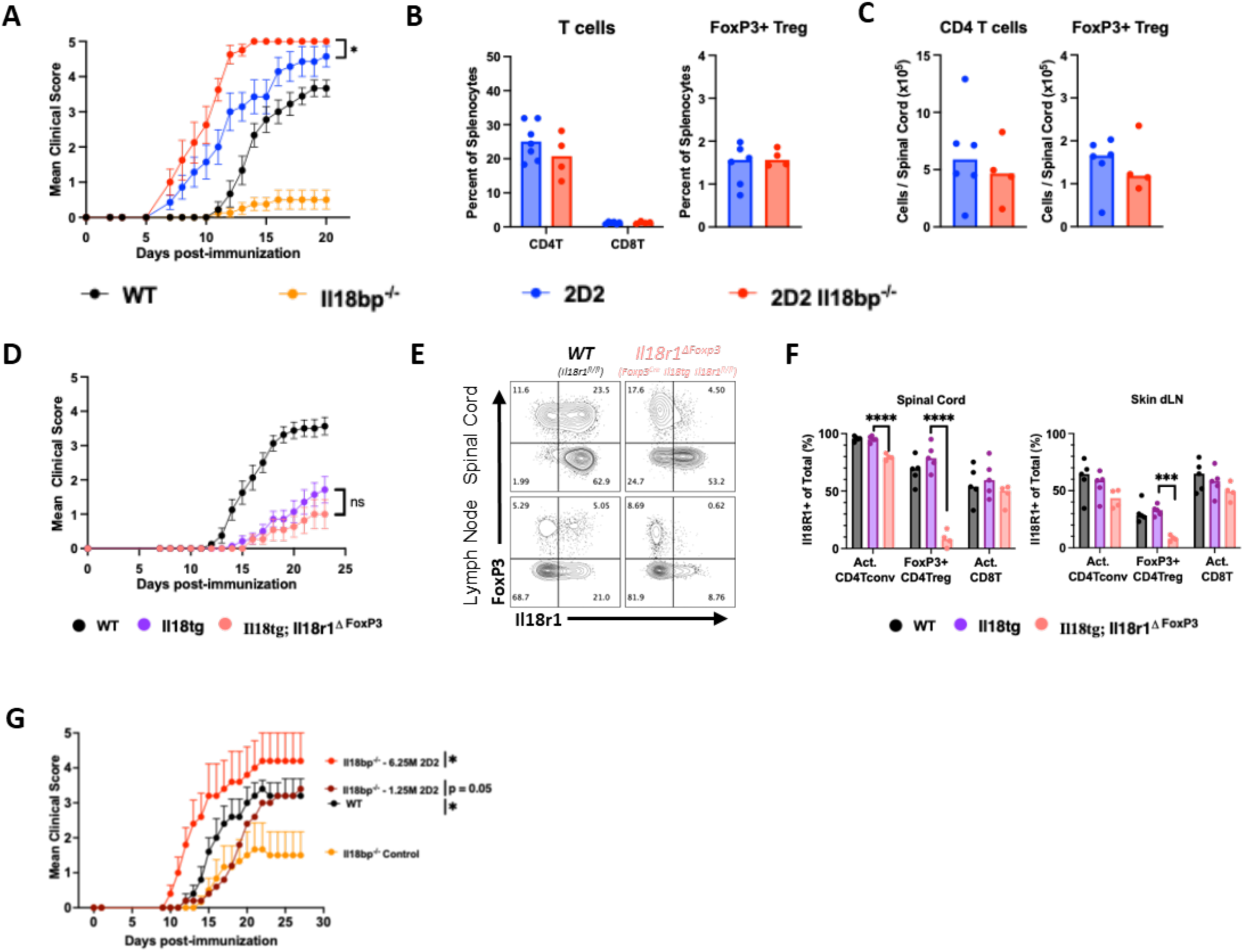
IL-18 does not directly inhibit autoreactive CD4 T-cells or protect through FOXP3^+^ CD4T_reg_. **(A)** Mean EAE clinical score of WT (n=9), *Il18bp^KO^* (n=8), 2D2 (n=7), or 2D2;*Il18bp^KO^*(n=8) mice. **(B,C)** Flow cytometric quantification of splenic and spinal cord T-cells at day 15. **(D)** Mean EAE clinical score of WT (n=16), *Il18tg* (n=14)*, or Il18tg;Il18r ^ΔFox3p^* (n=11) mice. **(E)** Representative flow plots of IL-18R1 and FOXP3 expression on day 23 CD4 T-cells. **(F)** Percent IL-18R1^+^ on CD44^+^ CD4T_conv_, CD4T_reg_, and CD44^+^ CD8T-cells. **(G)** Mean EAE clinical score of WT (n=4) or *Il18bp^KO^*(control n=6, 1.25M n=5,6.25M n=5) mice that received different amounts of splenic CD4 T-cells from naïve 2D2 mice on day −1. (A-D) Pooled data from 2 experiments. (E,F,G) Data representative of 2 experiments. Error bars = SEM. Statistical analysis: (A,D,G) Kruskal-Wallis of AUC with Dunn’s post-test of pairwise comparisons to *2D2;Il18bp^KO^* (A) or *ll18tg* (D) or *l18bp^KO^* control (G), (B) unpaired t-tests with Holm-Sidak correction for p-value, (C) unpaired t-tests, (F) one-way ANOVA with Dunnett’s post-test of pairwise comparisons to *Il18tg*. *p< 0.05, **p<0.01, ***p<0.001.

Intrigued by the paradoxical exacerbation of EAE in *2D2;Il18bp^KO^* mice, we wondered whether a surplus of autoreactive 2D2 T-cells would overcome protection in *Il18bp^KO^* mice. Transfer of 250,000 2D2 T-cells did not overcome protection (**Figure S3C**). *Il18bp^KO^* mice who received 1.25 million 2D2 T-cells prior to EAE induction still demonstrated a delay in EAE onset but equivalent peak clinical disease compared to WT. Transfer of 6.25 million 2D2s into *Il18bp^KO^* mice finally overcame protection by excess IL-18 (**Figure 4G).** Notably, we derived 2D2 T-cells from the spleens of unstimulated mice, and of these about 4% expressed FOXP3^+^. However, following transferring and EAE induction, < 2% of transferred 2D2 T-cells expressed FOXP3 (**DNS**). Thus, IL-18’s protection was resistant to transfer of >1 million MOG-autoreactive CD4 T-cells, and it did not appear to be mediated directly through CD4T_auto_ or CD4T_reg_.

### Excess IL-18 activates and expands effector CD8 T-cells, while CD8 depletion limits protection

In models of hyperinflammation, we have previously found that excess IL-18 preferentially activated CD8 over CD4 T-cells (16, 17). As such, and in light of our findings in EAE, we next considered whether CD8 T-cells were mediating IL-18’s protective activity. CD8 T-cells have been shown to play an important regulatory role in EAE and similar models of autoimmunity, but markers used to define a regulatory CD8 T-cell (CD8T_reg_) population have varied, including general effector/exhaustion markers like PD-1 and CD39, or more specific markers like CD122, Ly49, HELIOS, FOXP3, and/or CD38 (55, 56).

At day 12, splenic CD8 T-cells in *Il18bp^KO^* mice showed greater effector (CD44^+^CD62L^−^) than central-memory (T_cm_, CD44^+^CD62L^+^) activation, as well as higher expression of CD39 and PD-1 (**Figure 5A-B**). We have previously observed paradoxical IL-18R1 downregulation in *Il18tg* mice, likely as a regulatory response (16, 17). This heightened CD8 T-cell activation contrasted with reduced CD4 T-cell activation observed in *Il18bp^KO^* mice (**Figure 2**). Expression of other putative CD8T_reg_ surface markers was mixed: CD122 and Ly49 were reduced, FOXP3 expression was negligible in both genotypes, HELIOS (the transcription factor associated with Ly49+ CD8Treg (57)) was slightly elevated, and CD38 was increased (**Figure 5B & DNS**). In some circumstances, protection by CD8 T-cells correlated with IFNg production (58). Despite IL-18’s well-established effect on CD8 T-cell IFNg production, *ex vivo* IFNg production did not differ dramatically overall, with a trend toward greater PMA-induced IFNg production by WT and greater MOG-peptide-induced IFNg production by *Il18bp^KO^*CD8 T-cells (**Figure 5C**). Given our use of genetic models, it remained possible that these differences were unrelated to EAE induction. Without stimulation, CD8 T-cell phenotypes were remarkably similar between WT and *Il18bp^KO^* mice, whereas *Il18tg* mice (reported to have basal activation (15, 16)) showed modest, *IL18R1*-dependent changes specifically in HELIOS and Ly49 expression **(Figure S4A-B).** In the CNS, total CD8 T-cells were comparable between genotypes, despite greatly reduced CNS CD4 T-cells in *Il18bp^KO^*mice (**Figure 2D**, **Figure 5D-E)**. Differences in surface marker expression were blunted, but otherwise conserved between splenic and CNS CD8 T-cells (**Figure 5F**). A population of T-cells (CD44^hi^CD62L^−^) co-expressing CD38 and PD-1 was expanded in *Il18bp^KO^* mice in both spleen and PB at peak disease and seen in the CNS of all mice (**Figure 5G).** Thus, CD8 T-cell phenotyping suggests that excess IL-18 drives a program indicative of a highly-activated effector state.

**Figure 5.**
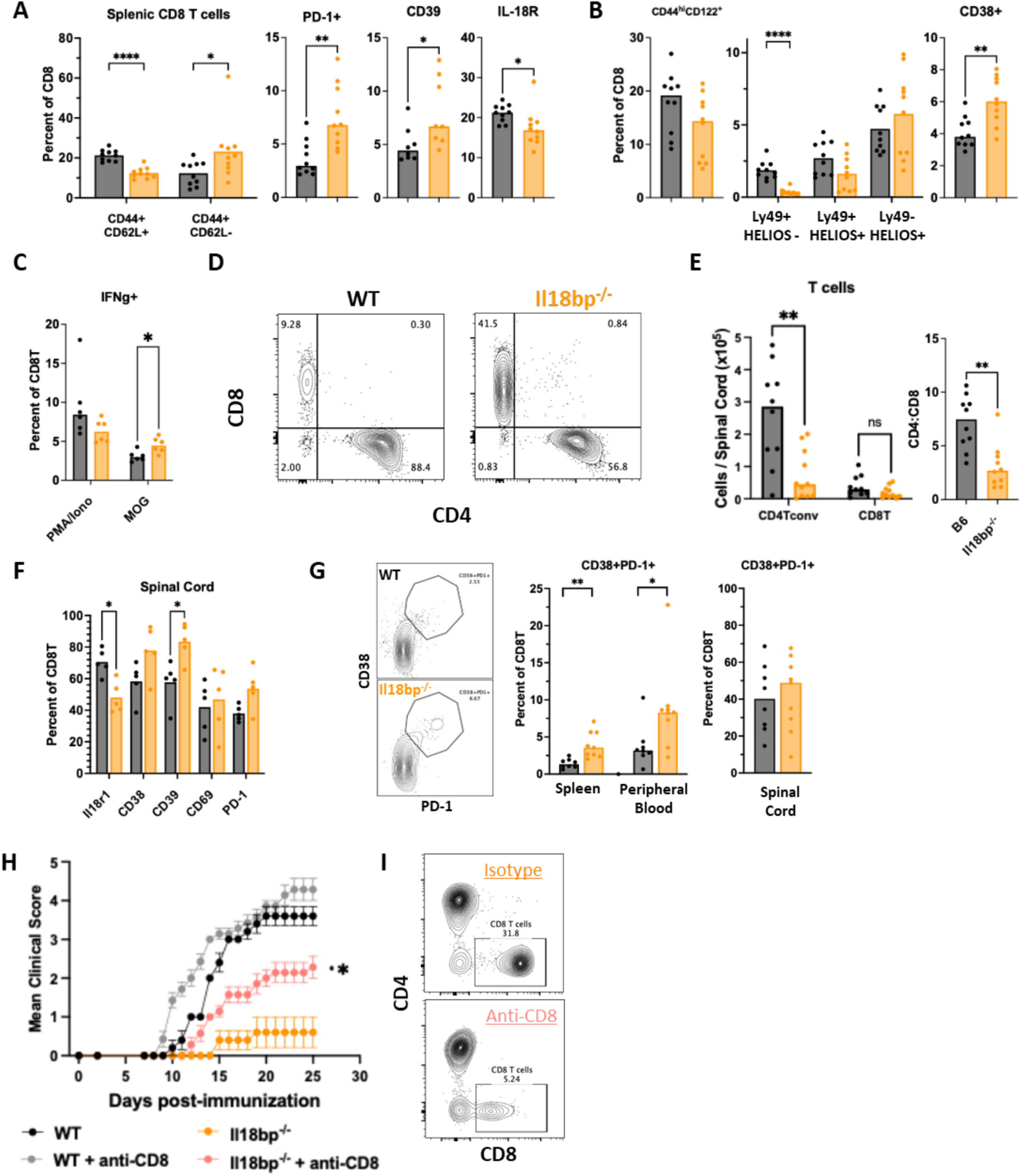
Excess IL-18 activates and expands effector CD8 T-cells, while CD8 depletion limits protection: Expression of **(A)** effector markers and **(B)** putative suppressive markers on splenic CD8 T-cells from WT and *Il18bp^KO^* mice at day 12. **(C)** IFNg production by day 12 splenic CD8 T-cells following PMA/Ionomycin or MOG^35-55^ stimulation by ICS. **(D)** Representative flow plots of day 18 spinal cord T-cells. **(E)** Quantification of spinal cord CD4 and CD8 T-cells and the corresponding CD4:CD8 ratio. **(F)** Expression of key effector surface markers on spinal cord CD8 T-cells. **(G)** Representative flow plots and quantification of dual-expressing CD38^+^PD-1^+^ CD8 T-cells. **(H)** Mean EAE clinical score of WT and *Il18bp^KO^*mice receiving isotype (n=5) or CD8-depleting antibodies (n=7; 200ug, i.p., days −3, 0, 3, and 6) (**I)** Representative plots of CD4 versus CD8 expression on splenic T-cells at day 25. (A,B, E, G, H) Data pooled from 2 experiments. (C,F) Data representative of 2-3 experiments. Error bars = SEM. Statistical analysis: (A-C, E-G) unpaired t-tests, p-value with Holm-Sidak correction with multiple comparisons, (H) Kruskal-Wallis of AUC with Dunn’s post-test of pairwise comparisons of *Il18bp^KO^* isotype vs anti-CD8. Only p_adj_<0.05 is shown. *p< 0.05, **p<0.01, ****p<0.0001.

We next examined their relevance by broadly depleting CD8 T-cells beginning at the time of EAE induction. Depleted WT mice showed a trend toward earlier disease onset, but *Il18bp^KO^* mice showed a more substantial loss but remained somewhat protected (**Figure 5H)**. Consistent with prior reports however (59), depletion of CD8 T-cells was incomplete in both spleens and draining lymph nodes and the remaining CD8T-cells contained a significant effector population (**Figure 5I and Figure S5A-B**)

### Excess IL-18 protects against EAE via IL-18-responsive CD8 T-cells and IFNg

To determine the extent to which IL-18’s protective effect in EAE depended on direct IL-18 sensing by CD8 T-cells, we bred *Il18tg* mice with tamoxifen-inducible deletion of *Il18r1* specifically on mature CD8 T-cells (*E8i^cre/cre^Il18r1^fl/fl^Il18tg, or Il18r1^ΔCD8^*). We induced EAE in *Il18tgIl18r1^ΔCD8^* mice (and relevant controls) and, to limit effects on priming and disease penetrance (60), administered tamoxifen on days 4 & 6 post-EAE induction. *Il18tgIl18r1^ΔCD8^* mice lost all protection conferred by IL-18 overexpression (**Figure 6A)**. As expected, IL-18R expression was lost specifically, but incompletely, on dLN and spinal cord CD8 T-cells of *Il18tgIl18r1^ΔCD8^* mice (**Figure 6B, Figure S5C**). Correspondingly, *Il18tgIl18r1^ΔCD8^* mice showed restoration of both dLN CD4T_conv_ CD44 expression and spinal cord CD4T_conv_ accumulation **(Figure 6C-D)**. Likewise, the accumulation of CNS CD8 T-cells in *Il18tg* mice was lost with e8i-specific deletion of *Il18r1*, significantly altering the CD4:CD8 ratio (**Figure 6D)**. *Il18tgIl18r1^ΔCD8^* mice enabled a more careful dissection of acute versus chronic changes. HELIOS expression by CD8 T-cells was diminished with broad, constitutive *Il18r1* deletion (CD4^Cre^, **Figure S4C**), but remained upregulated in *Il18tgIl18r1^ΔCD8^* mice (**Figure 6E-F**), suggesting HELIOS upregulation is indirect and unnecessary for protection. By contrast, temporal, e8i-specific deletion of *Il18r1* was sufficient to prevent downregulation of CD122 and Ly49 (**Figure 6E)**. As in other systems, we did not observe substantial differences in the phenotype of CNS-infiltrating T-cells, nor in FOXP3+ CD4T_reg_ peripherally or in the CNS (**Figures 6F-G**). Overall, these experiments suggest a potent, dominant, protective effect of IL-18 signaling specifically through mature CD8 T-cells.

**Figure 6:**
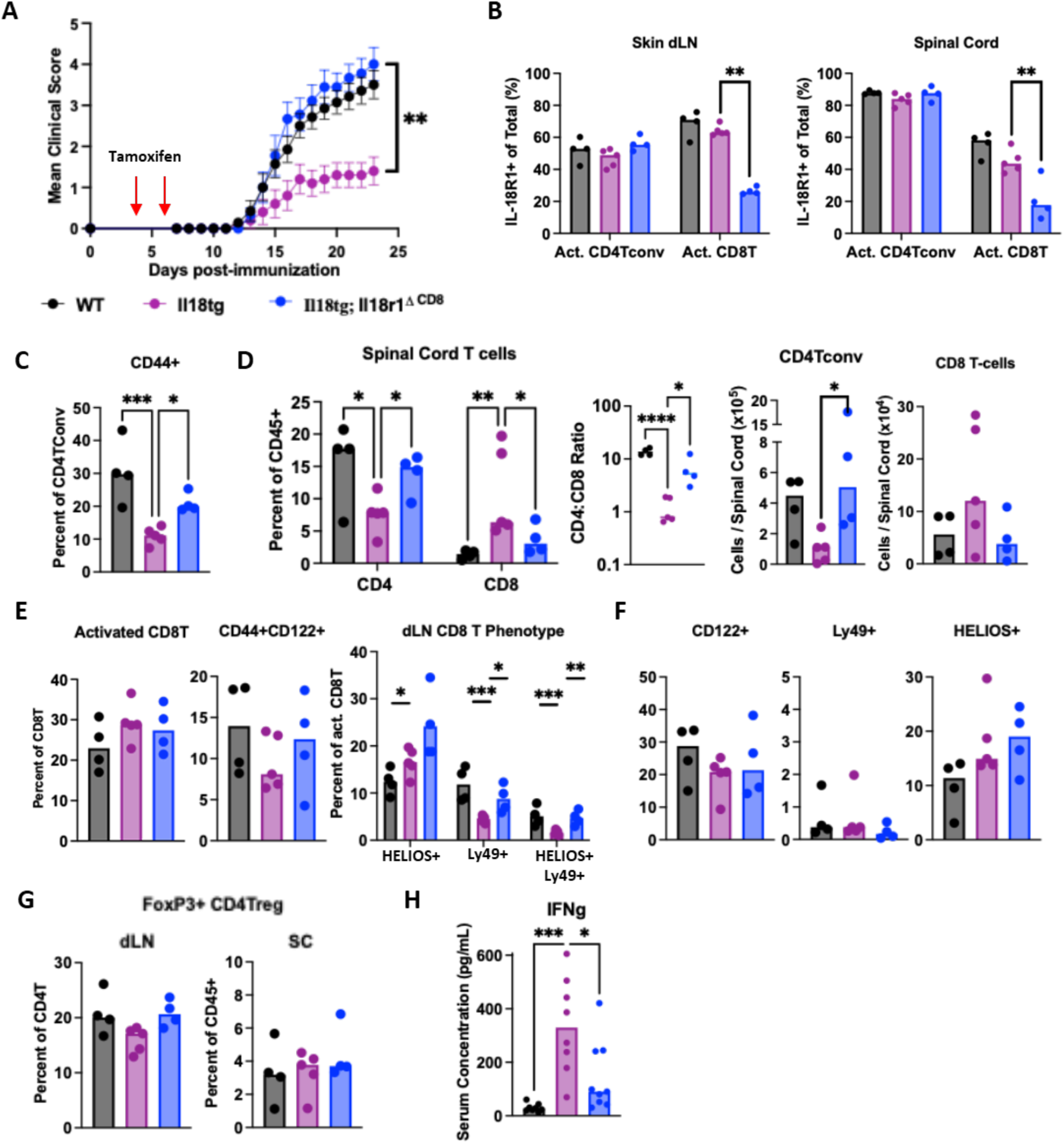
Excess IL-18 protects against EAE via IL-18-responsive CD8 T cells. **(A)** Mean clinical score of WT (n=12), *Il18tg* (n=10), and *Il18tg Il18r1^ΔCD8^* (n=9) mice treated with tamoxifen at days 4 and 6. **(B)** IL-18R1 expression on activated (CD44^+^) CD4Tconv and CD8 T-cells. **(C)** Expression of CD44 on CD4T_conv_ in dLN. (**D)** Quantification of T-cell subsets in day 23 spinal cords as percent of CD45, CD4:CD8 ratio, and absolute number. (**E)** CD8 T-cell activation (defined as CD62L-) and expression of putative CD8Tsupp markers in dLN. **(F**) CD8 T-cell expression of putative CD8Tsupp markers and (**G)** quantification of FOXP3+CD4T_reg_ in day 23 spinal cords. **(H)** Serum IFNg concentration of the indicated genotypes at day 23. (A) Data pooled from 2 experiments and representative of 3 total. (B-H) Data representative of 3 experiments. Error bars = SEM. Statistical analysis: (A) Kruskal-Wallis of AUC with Dunn’s post-test of pairwise comparisons, (B-H) one-way ANOVA with Dunnett’s post-test of pairwise comparisons, (D, E, G) 2way ANOVA with Dunnett’s post-test of pairwise comparisons. All comparisons made to *Il18tg* control. Only p_adj_<0.05 is shown. *p< 0.05, **p<0.01, ***p<0.001, ****p<0.0001.

CD8 T-cell IFNg production was modestly elevated in *Il18tg* mice and minimally affected in *Il18bp^KO^* mice (**Figures 3C & S4D**), and serum IFNg levels were likewise only modestly elevated (**Figures 1F & 6H**). Nevertheless, given their strong link we next assessed the role of IFNg in *Il18bp^KO^*mice. Consistent with prior reports (61, 62), continuous IFNg neutralization resulted in more severe EAE in WT mice. IFNg-blocked *Il18bp^KO^* mice were not only susceptible to EAE but their disease was comparable to IFNg-blocked WT mice (**Figure S6A).** IFNg also appears to be necessary for *in vitro*, contact-dependent suppression of CD4T_auto_ by CD38^+^ CD8 T-cells (58), and these data suggest IFNg plays a crucial role *in vivo* during IL-18-mediated protection from EAE as well. IFNg is a strong inducer of PD-L1 on APCs and other cells, and PD-L1 has a known suppressive effect on EAE (63, 64). However, PD-L1 blockade had no effect on protection in *Il18bp^KO^*mice (**Figure S6B**).

### DR-18, synthetic IL-18 with engineered resistance to IL-18BP, disrupts autoimmune effector activity and protects from EAE immunopathology

Targeting specific cytokines has been a transformative therapeutic strategy in dozens of inflammatory diseases. Given profound protection by excess IL-18 in genetic systems, we wondered whether exogenous IL-18 could be effective by itself or as part of immunoregulatory CD8 T-cell therapies. Adapting an established strategy (65, 66), we cultured CD8 T-cells (from day 10 EAE spleens) with MOG^35-55^ peptide +/− IL-18 for 3 days before transferring into WT mice and inducing EAE. CD8 T-cell transfer was not sufficient to prevent EAE, but IL-18-stimulation of CD8 T-cells prior to transfer substantially delayed onset relative to controls (**Figure S6A)**, suggesting that IL-18’s protection required more continuous exposure. However, recombinant murine IL-18 given every other day did not significantly protect WT mice (**Figure S6D)**, possibly because we failed to overcome neutralization by endogenous IL-18BP.

DR-18 proteins are a class of synthetic IL-18 analogs selected for their resistance to neutralization by IL-18BP and retained IL-18R signaling (24). The human DR-18 ST-067 is currently under clinical investigation in cancer immunotherapy. DR-18 conferred near-complete protection to WT mice from paralysis and weight loss (**Figures 7A & S6E)**. As with *Il18tg* mice, protection by DR-18 was largely dependent on CD8 T-cell expression of *Il18r1* (**Figure 7B)**. An exogenous system of protective IL-18 enabled us to interrogate granule-mediated cytotoxicity, which appears required for CD8 T-cell protection from EAE in some contexts (65, 67). Perforin-deficiency did not appreciably exacerbate EAE in WT mice, nor did it significantly affect protection by DR-18 (**Figure S6F**). Therapeutic DR-18 also enabled an examination of the timing of protection. We treated WT mice with DR-18 at various dosing schedules and found the most profound protection occurred with as little as two doses administered as late as day 12 (**Figure 7C**), well after CD4T_auto_ induction and substantial expansion.

**Figure 7.**
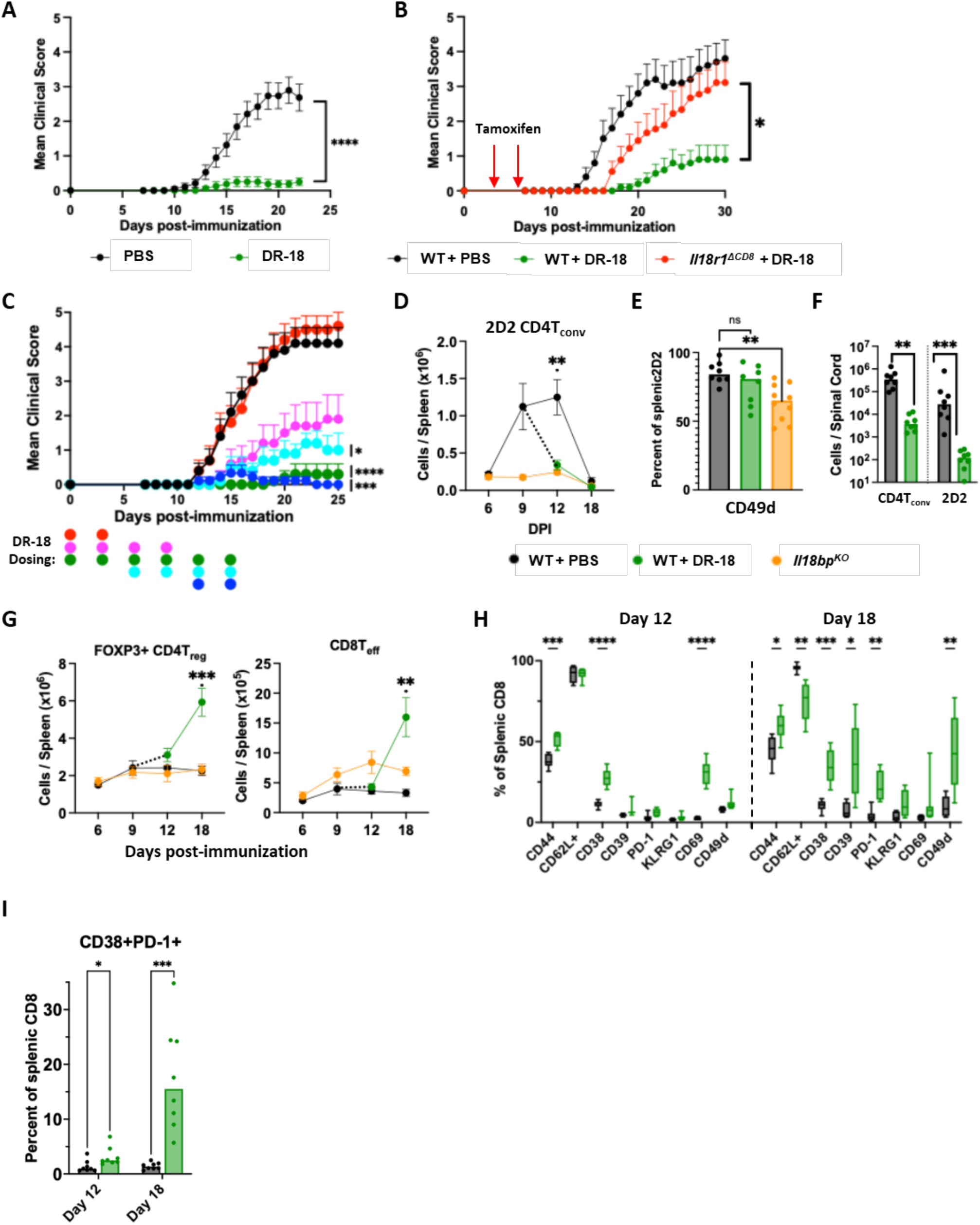
DR-18 disrupts autoimmune effector activity and protects from EAE immunopathology. **(A)** Mean EAE clinical score of WT mice treated with PBS (n=19) or DR-18 (2ug, Subq, q3d; n=16) from days 0 to 15. **(B)** Mean EAE clinical score of WT and *E8i^cre^;Il18r1^fl/fl^*mice treated with PBS or DR-18 (n=9; 2ug, treated every 3 days from 0 to 30). **(C)** Mean EAE clinical score of WT mice treated with PBS or DR-18 at various timepoints indicated below the graph (treatment every 3 days from 0 to 15, colored dot corresponds to DR-18 treatment while no dot indicates PBS; n=10 for all groups). **(D-I)** 2.5×10^5^ splenic CD4 T-cells from naïve 2D2 mice were transferred to WT and *Il18bp^KO^* mice on day −1. Half of the WT mice were randomized to receive DR-18 (n=8 per timepoint) or PBS (n=8 per timepoint) on days 9, 12, & 15. **(D)** Quantification of transferred splenic 2D2 T-cells, **(E)** CD49d expression on day 12 splenic 2D2, **(F)** and CD4T_conv_ and 2D2 in day 18 spinal cord. **(G)** Absolute number of splenic FOXP3+ CD4T_reg_ and CD8T_eff_ (CD44^+^CD62L^−^) cells over time. **(H)** Expression of various surface markers and **(I)** CD38/PD-1 co-expression on splenic CD8 T-cells at day 12 and 18 from WT mice treated with PBS and DR-18. Data pooled from (B-I) 2 or (A) 3 experiments. Error bars = SEM except for box and whisker plot. Statistical analysis: (A) Mann-Whitney of AUC, (B,C) Kruskal-Wallis of AUC with Dunn’s post-test of pairwise comparisons to DR-18 (B) or PBS control (C), (D, G) unpaired t-tests of WT vs WT+DR-18 on days 12 and 18 only with Holm-Sidak correction of p-value, (E) one-way ANOVA with Dunnett’s post-test of pairwise comparisons (H,I) unpaired t-tests with Holm-Sidak correction of p-value. Only p_adj_<0.05 is shown. *p< 0.05, **p<0.01, ***p<0.001, ****p<0.0001.

The ability of DR-18 to protect within such a short window enabled a careful, temporal examination of its effects on competing T-cell populations. Since we observed robust CD4T_auto_ activation and expansion in WT mice by day 12 (**Figure 2),** we transferred 2D2 T-cells prior to EAE induction and then randomized WT mice to receive DR-18 or PBS. DR-18 treatment from day 9 to 15 completely protected from EAE (**Figure S7A**). At day 6, WT and *Il18bp^KO^* mice had similar numbers of 2D2 CD4T_auto_ in both spleen and peripheral blood. These 2D2 cells continued to expand in the periphery through day 12, yet they stagnated in *Il18bp^KO^* mice after day 6 (**Figures 7D and S7B,C**). By day 12, the number of transferred 2D2 T-cells was dramatically reduced in the spleen and PB of WT mice with only three days of DR-18 exposure. Peripheral 2D2 cells declined in all mice by day 18. Unlike our observations in *l18bp^KO^*mice, we did not see a consistent decrease in CD49d-expressing 2D2 T-cells at day 12, reinforcing the findings that IL-18 can suppress highly activated effector CD4T_auto_ **(Figure 7E).** However, consistent with their low clinical scores, DR-18-treated and *Il18bp^KO^*mice demonstrated very few spinal cord CD4 T-cells (2D2 or endogenous) at the peak of clinical symptoms (**Figure 7F**). DR-18 treatment led to the accumulation of both FOXP3^+^ CD4Tregs and effector CD8T-cells greater than observed in control mice (**Figure 7G, S7D)**. The accumulation of effector CD8 T-cells was associated with profound peripheral effector CD8 T-cell activation through days 12 and 18. On day 12, CD8 T-cells from DR-18 mice demonstrate increased CD44, CD38, and CD69-expressing cells (potentially indicative of early activation). Nearly all surface markers associated with activation are increased on DR-18 treated CD8 T-cells by day 18 except for CD69. Activation of the CD8 T-cell compartment is best seen by the substantial increase in the dual-expressing CD38^+^PD-1^+^ population previously seen in *Il18bp^KO^* mice (**Figure 7H, S7E-F**). This population accumulated more substantially in *Il18bp^KO^*than in WT mice, and their growth was explosive following DR-18 treatment (**Figure S7F).** Thus, acute DR-18 administration was dramatically protective, with an acute reciprocal shrinking of autoreactive T-cell populations and expansion of effector CD8 T-cells. Overall, these data support a model wherein excess IL-18 drives an effector response in CD8T-cells and subsequent and specific IFNg-dependent suppression of autoreactive CD4 T-cells, preventing accumulation of CNS-toxic T-cells, histologic white matter damage, and clinical deterioration.

## DISCUSSION

For autoreactive T-cell clones to drive autoimmunity, they must encounter self-antigens in the presence of co-stimulatory signals. IL-18 promotes co-stimulation both indirectly, by inducing cytokines such as IFNg that act on antigen-presenting cells, and directly on non-naïve T-cells. Thus, diseases of chronic IL-18 elevation illustrate an important immunologic paradox: why doesn’t chronic innate immune activation *always* lead to autoimmunity? Regulatory T-cells are an important, but poorly understood, part of the answer. Foxp3^+^ CD4T_reg_ are well-recognized for this ability, but they may be most efficacious in limiting humoral autoimmunity (68). Opposing cellular autoimmunity, particularly the direct pathogenic effects of CD4T_auto,_ may become increasingly relevant in a future of deep B-cell depletion (69). Although controversy around “T-suppressor” cells discouraged studies of regulatory CD8 T-cells (70), recent discoveries have revitalized interest (56).

We evaluated this paradox in EAE, a model useful for preclinical MS studies and for studying *in vivo* competition between pathogenic CD4T_auto_ and multiple populations of regulatory T-cells. Prior EAE studies suggested a pathogenic role for IL-18: IL-18 deficiency and CNS delivery of IL-18BP modestly attenuated disease (36, 40, 41). Even without extra IL-18, pathogenic CD4T_auto_ secrete canonical IL-18-induced cytokines like IFNg and GM-CSF (42, 43, 47), and CD4T_auto_ stimulated *in vitro* with IL-18 drive more severe EAE upon transfer (71, 72). Thus, we expected that excess IL-18 would act on the dominant IL-18 responsive T-cell in EAE, CD4T_auto_, and exacerbate CNS inflammation.

By contrast, excess IL-18 was profoundly protective across three distinct systems and two mouse colonies, opposing CD4 autoreactivity and tipping the scales against autoimmunity. Exogenous IL-18 may also be therapeutic in the NOD model of Type 1 diabetes (73). We observed neither disease improvement nor worsening in *Il18^KO^* mice (**Figure S1**) and found that the effects of excess IL-18 on CD8 T-cells readily outweighed its modest amplification of CD4T_auto_ (observed only in 2D2 mice, **Figure 4A**).

Excess IL-18’s proclivity for CD8 over CD4 T-cell activation has been observed in other contexts. In models of HLH involving excess IL-18 and a variety of triggers, CD8 T-cell activation far outstripped that of CD4s (13–17). In tumor immunotherapy, DR-18 drove enhanced CD8 T-effectors and stem-like precursors to mediate tumor control (24). IL-18’s preference for CD8’s is not absolute, as observed in 2D2 mice and in *T. gondii* infection, where antigen-specific CD4 T-cell responses are particularly robust (74).

Though regulatory CD8 T-cells are critical for preventing immunopathology in EAE and beyond (56), defining CD8T_reg_ has been complicated without a defining marker like “Foxp3”. We sought to characterize IL-18-activated CD8 T-cells in EAE by *phenotype*, *antigen specificity*, and *effector functions*. Among candidate CD8T_reg_ phenotypic markers, only CD38 and Helios increased consistently with excess IL-18, and Helios was not sufficient for protection (**Figure 4F**). Prior studies of protective CD38^+^CD8 T-cells identified a central-memory (T_cm_-like) phenotype characterized by expression of CD44, CD122, and CD62L (58). By contrast, we found CD8 T-cells exposed to excess IL-18 looked most like CD8 T_effector_ cells, with low expression of CD62L and upregulation of KLRG1, PD-1, CD39, CD49d, and CD11a. IL-18 is a potent driver of the effector T-cell program, so one explanation for this effector phenotype is that canonical (T_cm_-like) CD8T_reg_ adopted CD8T_effector_ features upon exposure to IL-18. Alternatively, IL-18 may protect via CD8 T-cells arising distinctly from traditional CD8T_reg_. Heterogeneity in a “CD8T_reg_ phenotype” is not without precedent: the CD8 T-cell exhaustion program varies substantially between tumors, chronic viral infections, or in hyperinflammation (17, 75). However, the various flavors of exhausted CD8 T-cells share a core surface and transcription factor signature, and such a core signature has not yet been detected in CD8T_reg_ studies.

CD8T_reg_ may be better defined by MHC-I and antigen specificity, and TCR signaling is a prerequisite for sustained expression of the IL-18 receptor (17). Murine CD8T_reg_ have been described to use either classical (65, 67) or non-classical MHC-I (76–79), and recognize antigens derived from autoreactive cells themselves (79), neuropeptides (65, 80, 81), or unidentified antigens (67). Notably, relapsing MS patients may have a deficit in neuropeptide-specific CD8T_reg_ (82, 83). In MAS models, IL-18 promoted T-cell clonality but did not favor specific clones between mice (16, 17). In *ex vivo*-stimulations, splenic CD8 T-cells from *Il18bp^KO^* mice showed greater cytokine production with MOG-peptide stimulation, but this difference was lost with more broad stimuli (**Figure 5C)**. This suggests that the CD8 T-cells critical for protection respond to either MOG-peptide themselves and/or to adjacent MOG-responsive cells (likely CD4T_auto_). Nevertheless, it remains unclear whether excess IL-18 alters the nature of antigen presentation to protective CD8 T-cells in EAE.

Like their more famous FoxP3+ counterparts, CD8T_reg_ have been shown to protect via multiple mechanisms including contact-dependent cytotoxicity (both Granzyme- and FasL-mediated) and regulatory cytokine secretion (58, 76, 84–86). We found that neither perforin nor PD-L1 signaling were necessary for IL-18 to protect. However, IFNg neutralization abrogated protection in *Il18bp^KO^* mice, despite relatively meager increases in systemic IFNg levels (**Figures 1F**, **6H**) (14). These findings parallel others’ observations that CD38^HI^ CD8T_reg_ require both cell contact and IFNg to suppress CD4 T-cells (58, 87). Overall, our data links IL-18 to the biology of CD8T_reg_ cells but their collective identity remains unclear.

The phases of autoimmune development have been well-characterized in EAE, and IL-18’s protection was notably late in the process. DR-18 treatment at only early timepoints may have mildly promoted EAE, consistent with prior data suggesting an early pathogenic role (40). Likewise, excess IL-18’s protective effect on transferred autoreactive 2D2 cells was not observed until after priming and early expansion (∼days 9-12). Critically, exposure to excess IL-18 at this late phase was both necessary and sufficient for protection: Loss of IL-18 responsiveness, even after a lifetime of chronic IL-18 exposure, abrogated protection (**Figure 6**). DR-18 treatment at only this late expansion phase was as protective as continuous dosing. Similar biphasic effects have been reported for other Type 1 cytokines (88–91).

Geographically, excess IL-18 diminished CD4T_auto_ in the periphery and reduced accumulation of any white blood cells in the spinal cord. CNS CD4 and CD8 T-cells were phenotypically similar, but far less abundant, in mice with excess IL-18 than WT. This correlated with specific reduction of peripheral CD4T_auto_ that expressed the CNS-homing integrin and MS therapeutic target CD49d (92). Together, these findings support a model where excess IL-18, via CD8 T-cells, targets highly-activated CD4T_auto_ before they enter the CNS and potentially also after their arrival.

Our results warrant both deeper and broader preclinical validation. MOG^35-55^-induced EAE is a useful autoimmune model, but incompletely reflects the relapsing-remitting course typical of MS. Our studies also do not address how excess IL-18 affects the autoreactive T-follicular/helper cells (Tfh) or B-cells relevant in other autoimmune models. Validation in human cells/systems is also necessary prior to consideration of IL-18 for use in human autoimmunity.

MS is the best example of an autoimmune disease successfully treated with cytokine therapy, namely IFNβ (93, 94). Our and others’ data suggest other Type 1 cytokines may also have a similar effect on EAE, although the specificity of IL-12, IL-15, or IFNg for CD8T_reg_ populations has not been established (55, 79, 95). A small trial of systemic IFNg in MS was not successful (96). Notably, the MS therapy glatiramer acetate induces CD8T_reg_ as part of its mechanism, supporting the therapeutic relevance of this pathway (84). Even if effective, safely harnessing protection by IL-18 will require a better understanding of how to avoid provoking MAS or amplifying autoreactive responses (as observed in 2D2 mice). We did not observe MAS or dramatic increases in systemic IFNg levels with EAE induction relative to TLR9 stimulation or LCMV infection (13, 16), but such control over environmental stimuli is not practical in patients. Nevertheless, IL-18 elevation is not sufficient for disease activity in NLRC4-GOF patients (97) and IL-18-based immunotherapies currently in cancer trials (including DR-18) appear to be well tolerated relative to other immunostimulatory cytokines (98). Using cytokines to prepare regulatory cell therapies, such as low-dose IL-2 to expand CD4T_reg_ (99), may avoid these systemic risks altogether (100, 101).

Intracellular bacterial infections (e.g. *Y. pestis*) once drove evolutionary selection for potent inflammasome responses (102). In EAE, an inflammatory model where autoreactive and regulatory T-cells compete, we found an unexpected role for inflammasome-activated IL-18 in tipping the scales toward regulation. It did so through preferential effects on CD8 T-cells and potent amplification of effector programs – including IFNg. These are the same mechanisms by which IL-18 appears to drive MAS and that inspired its investigation as a cancer immunotherapy. This provocative extension of IL-18’s therapeutic potential, though grounded in the comparative study of human immune dysregulation, requires replication, validation, and translation. Nevertheless, these findings suggest a novel therapeutic strategy borne of the link between ancient infectious pressures and the need to regulate self-autoreactivity.

## METHODS

### Sex as a biological variable

All experiments were performed on male mice for consistent EAE induction with baseline protection experiments repeated in female mice.

### Mice

C57Bl/6 (664), 2D2 (6912), *Prf1^−/−^* (2407), *CD4^cre^* (22071) and *Foxp3^yfp-cre^* (16959) mice originated from Jackson Laboratories. *Il18bp^−/−^* mice were obtained from the Knockout Mouse Project*. Il18tg* mice were a gift from Tomoaki Hoshino (Kurume University), *Il18r1^fl/fl^* mice were a gift from G. Trinchieri (National Cancer Institute), *E8iCre^ERT2/GFP^* mice were a gift from D. Vignali (University of Pittsburgh).

### Experimental Autoimmune Encephalomyelitis Induction

EAE was induced in 8 to 14 week old mice by subcutaneous injection of 200uL of CFA/MOG^35-55^ emulsion on day 0 with 500 uL of pertussis (400-500ng based on reported potency, i.p, List labs) on days 0 and 2. Emulsions were prepared by 2-syringe method using equal parts 1mg/mL mouse MOG^35-55^ (Biosynthesis) and 10 mg/mL CFA (made from BD DIFCO IFA with BD DIFCO H37RA mycobacterium). Clinical score was evaluated daily after day 7 using an established scale from 0 to 5 as follows: 0 = no symptoms, 1 = tail limpness, 2 = tail limpness and hind limb weakness (presenting as wobbly gait or impaired righting reflex), 3 = hind limb paralysis, 4 = complete hind limb paralysis with forelimb weakness, 5 = moribund or death.

### In vivo treatment

Antibodies: Mice were treated with various monoclonal antibodies and appropriate isotype-control antibodies via i.p. injection as follows: Anti-IFNg antibody (200ug, XMG1.2, Tonbo Biosciences) on days 1, 4, and 8. Anti-PD-L1 (250ug, 10F.9G2, BioXcell) every 3 days from day 0 to 21. Anti-IL-18R1 antibody (500ug per dose, Clone 9E6, Genentech) at various timepoints indicated in the figure. Anti-CD8a depleting antibody (200ug, YTS169.4, BioXcell) on days −3, 0, 3, and 6.

Cytokines: Mice were treated with recombinant murine IL-18 (1ug, i.p, MBL Life Science.) every other day starting at either day 1 or day 9. Mice were treated with DR-18(24) (2ug, subcutaneous, CS2 variant, Simcha Therapeutics Inc.) every 3 days from day 0 to 15 unless otherwise indicated.

Tamoxifen: Mice were injected with tamoxifen (1mg, i.p,, SIGMA) suspended in sunflower oil (SIGMA) on days 4 and 6 post-EAE induction.

### 2D2 T-cell isolation and transfer

Splenocytes were harvested from naïve *2D2*;*Il18bp^KO^* mice via mechanical dissociation and CD4 T-cells were isolated by negative magnetic sorting via Mojosort mouse CD4 T-cell isolation kit (Biolegend). In some experiments, 2D2 T-cells were subsequently labeled with Tag-it-Violet proliferation (BioLegend) dye following manufacturers’ protocol. Cells were resuspended in PBS (Corning) and adoptively transferred at appropriate concentration via retroorbital injection. 2D2 T-cell survival and expansion was assessed by flow cytometry.

### CD8 T-cell isolation and transfer

WT mice were immunized with MOG^35-55^ for EAE induction and spleens were harvested on day 10. Splenocytes were cultured for 3 days with rmIL-2 (10pg/mL. Peprotech), MOG^35-55^ peptide (20ug/mL, Biosynthesis), +/− IL-18 (50pg/mL, MBL Life Science). After 72 hours, CD8 T-cells were isolated by magnetic sorting using MojoSort Mouse CD8 T Cell Isolation Kit (Biolegend8). 5×10^6^ CD8 T-cells were transferred to naive WT mice by retro-orbital injection followed by EAE induction 24 hours later.

### Tissue collection and cell isolation

Leukocytes from spleen and draining lymph node were isolated by mechanical dissociation through a 100uM strainer. Spinal cord mononuclear cell isolation was adapted from a published protocol (103). Briefly, mice were perfused with cold PBS (Corning) by cardiac puncture and spinal cords were extruded into cold RPMI (Cytiva). Cords underwent mechanical and enzymatic digestion (collagenase and DNAse I in RPMI, 37c for 30 min) followed by 30/70% Percoll (Cytiva) density gradient separation and leukocytes were collected from the interphase for further analysis. Whole blood was collected into EDTA tubes and complete blood counts were assessed using a Drew Scientific Hemavet 950. For flow cytometry, whole blood was lysed with ACK lysis buffer (Quality Biological) and filtered through a 100uM strainer. Serum was isolated from whole blood by centrifugation using standard serum separator tubes. All single-cell suspensions were prepared in PBS with 2% fetal-bovine serum (Atlanta Biologicals) for flow cytometry.

### Flow cytometry and analysis

Single cell suspensions were stained with surface antibodies in HBSS (GIBCO) for 30 minutes followed by fixation and intracellular staining using eBioscience FOXP3/Transcription factor staining buffer set (Thermofisher). Samples were acquired using a 5-laser Cytek Aurora spectral cytometry or Beckman Coulter Cytoflex LX cytometer and analyzed on FlowJo v10.10.

For leukocyte analysis, cells were defined as follows: splenic myeloid (NK1.1^−^CD11b^+^), NK cells (B220^−^/TCRb^−^/NK1.1^+^), T cells (NK1.1^−^/CD11b^−^/B220^−^/TCRb^+^/CD4^+^ or CD8^+^). T-cells were further divided as follows: CD4T_conv_ (FOXP3^−^), CD4T_reg_ (FOXP3^+^), 2D2 (CD4^+^CD8^−^/TCRVa3.2^+^/TCRVb11^+^; **Fig S3A,D**), T_eff_ (CD44^hi^CD62L^−^), central memory T/T_cm_ (CD44^hi^/CD62L^+^), and naive T (CD44^lo^,CD62L^−^). Spinal cord leukocytes were assessed using the same criteria except all leukocytes were initially defined by CD45 expression and myeloid populations were subdivided between CD45^hi^ and CD45^mid^ **(Fig S2F)**. For UMAP visualization of day 12 splenic samples, an equal number of WT and *Il18bp^KO^* CD4 T-cells were concatenated from 4 samples per group. UMAP visualization was performed using FlowJo Exchange UMAP plugin (v4.1.1) only on 2D2 T-cells in the concatenated samples.

### *Ex vivo* T cell stimulation

Cells were plated at 2M/mL in R10 media (RPMI1640; GIBCO, 10% FBS; Atlanta Biologicals) and incubated for 5 hours with PMA/Ionomycin/Brefeldin A (BioLegend, cell activation cocktail with Brefeldin A) or 18 hours with MOG^35-55^ peptide or no peptide. For peptide stimulation, Brefeldin A (BioLegend) was added for the final 5 hours of culture followed by intracellular staining as above. For cytokine measurements in supernatant, cells were isolated and plated in the same conditions above for 72 hours with MOG^35-55^ without brefeldin A.

### Cytokine measurements

IFNg serum levels were assessed by BD OptEIA Mouse IFN-γ ELISA following manufacturers protocol (BD Biosciences) with absorbance measurements using ThermoFisher VarioSkan Lux. All other cytokine and chemokine measurements were assessed using mouse cytometric bead array kits (BD Biosciences) following manufacturers protocol. Samples were analyzed using 3-laser Beckman Coulter CytoFLEX Flow cytometer. All cytokine measurements shown are the average of 2 technical replicates with a baseline offset equal to the lowest standard (3.5pg/mL in ELISA, 5 pg/mL in CBA)

### Histology

Mice were perfused with PBS followed by 4% paraformaldehyde (PFA) via cardiac puncture. Spinal cords were gently extruded from the vertebral column using a PBS-filled syringe. Cords were immediately fixed for 48 hours in 4% PFA, processed using Tissue-Tek VIP 6AI Tissue processor, and paraffin embedded in a sagittal orientation. 5 micrometer paraffin-embedded sections were cut, deparaffinized, and stained with hematoxylin and eosin. Inflammatory foci were counted at 100x magnification for the length of intact cord per each section.

### Statistics

Frequentist statistical analyses were performed in GraphPad Prism v10 on representative or pooled experiments as indicated in figure legends. EAE experiments were analyzed by calculating individual area under the curve (AUC) of clinical disease score over time for each mouse followed by the appropriate statistical tests indicated to evaluate significance. Significance represented by number or symbol where ns = not significant, * = p<0.05, **=p<0.01, *** =p<0.001, ****=p<0.0001

### Study Approval

All animal studies were performed with approval from the Institutional Animal Care and Use Committee (IACUC) of The Children’s Hospital of Philadelphia or University of Pittbsurgh.

## Data availability

All data and materials will be made available upon request to corresponding author.

## Acknowledgments

The authors would like to thank Dr. Aaron Ring and Simcha Therapeutics, Inc., for providing murine DR-18, Dr. Chris Hunter and Dr. Mandy McGeachy for providing reagents, and Dr. Katherine Poholek for her experience with EAE experiments. We would also like to acknowledge the Flow Cytometry core at the Children’s Hospital of Philadelphia (CHOP) for their assistance.

## Funding

NIH/National Institute of Child Health and Human Development grant: R01HD098428 (SWC) RRF Rheumatology Future Physician Scientist Award: 1269102 (JAM)

## Conflict-of-interest

SC has received in-kind support (reagent) from Simcha Therapeutics, been site PI for a Novartis-sponsored trial, received support for speaking by Sobi & BMS, and been an ad hoc consultant for Apollo therapeutics, BMS, Johnson & Johnson, Novartis, and Sobi. SC, VD, and JAM are named in provisional patent PCT/US24/37468 “Methods for modulating activity of IL-18 for treatment of CD4+ T cell mediated autoimmunity and immunopathology.”

## Author contributions

Conceptualization: JAM, VD, LH, SWC

Investigation: JAM, VD, LH, JV, ZL, AFK, SWC

Visualization: JAM, VD, LH, AFK, SWC

Funding acquisition: SWC, JAM

Writing – original draft: JAM, SWC

Writing – review & editing: JAM, VD, LH, JV, ZL, AFK, SWC

