## Supplemental Figures 1-7 for "The innate cytokine IL-18 inhibits CNS autoimmunity through preferential activation of protective CD8 T-cells"

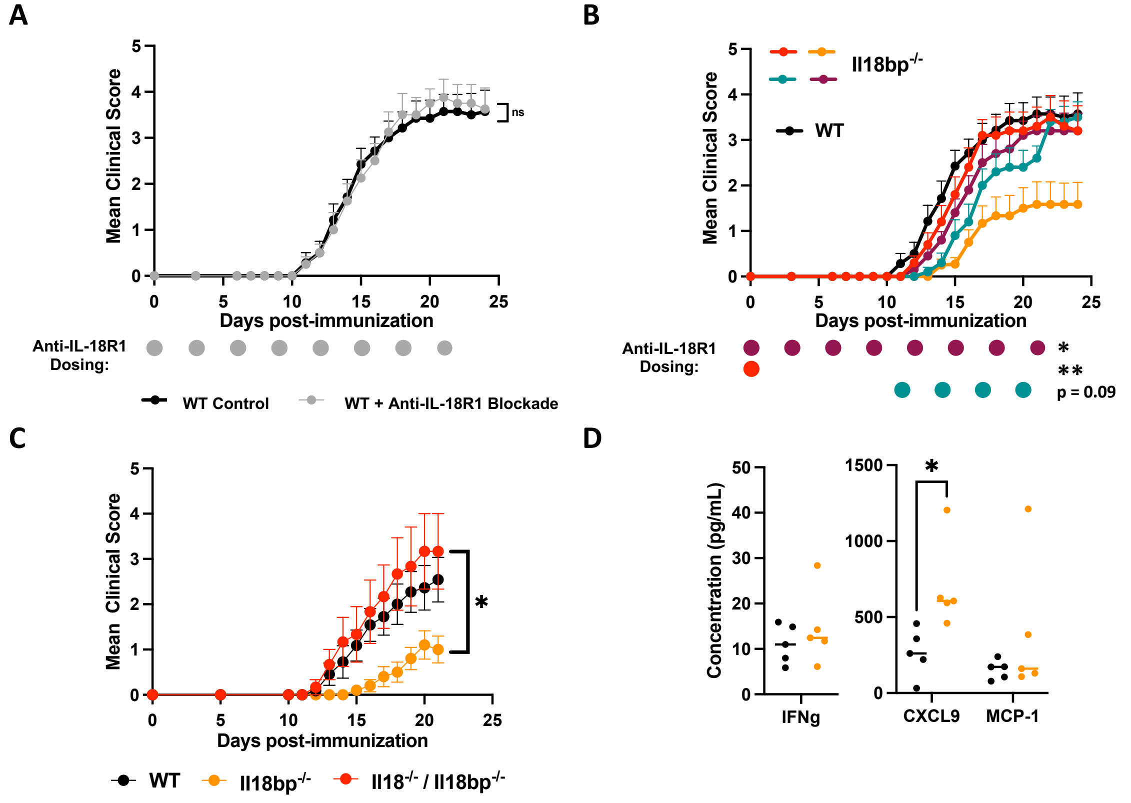


**Supplemental Figure 1: IL-18 protects against EAE through the canonical IL-18-receptor.** **(A)** Mean clinical scores following MOG$^{35-55}$ EAE induction of WT mice treated with 500ug anti-IL-18R1(every 3 days, day 0-21; n=8) or control (n=14). **(B)** Mean clinical score of WT, *Il18bp$^{KO}$* control (orange, n=12), or *Il18bp$^{KO}$* mice treated with **500ug anti-IL-18R1** at various time-points as indicated by the corresponding color dots below the graph (maroon n=10, red n=10, blue n=10). **(C)** Mean clinical score of WT (n=11), *Il18bp$^{KO}$* (n=10), and *Il18$^{KO}$;Il18bp$^{KO}$* (n=7) mice following MOG$^{35-55}$ EAE induction. **(D)** IFNg and IFNg-inducible chemokine levels from skin-draining lymph node homogenate at day 12.

(A-C) Data pooled from 2 experiments. (D) Data representative of 2 experiments. Error bars = SEM. Statistical analysis: (A) Mann-Whitney of AUC, (B, C) Kruskal-Wallis of AUC with Dunn's post-test of pairwise comparisons comparison made to *Il18bp$^{KO}$*, (D) unpaired t-tests, p-value with Holm-Sidak correction. ns = not significant, *p< 0.05, **p<0.01.

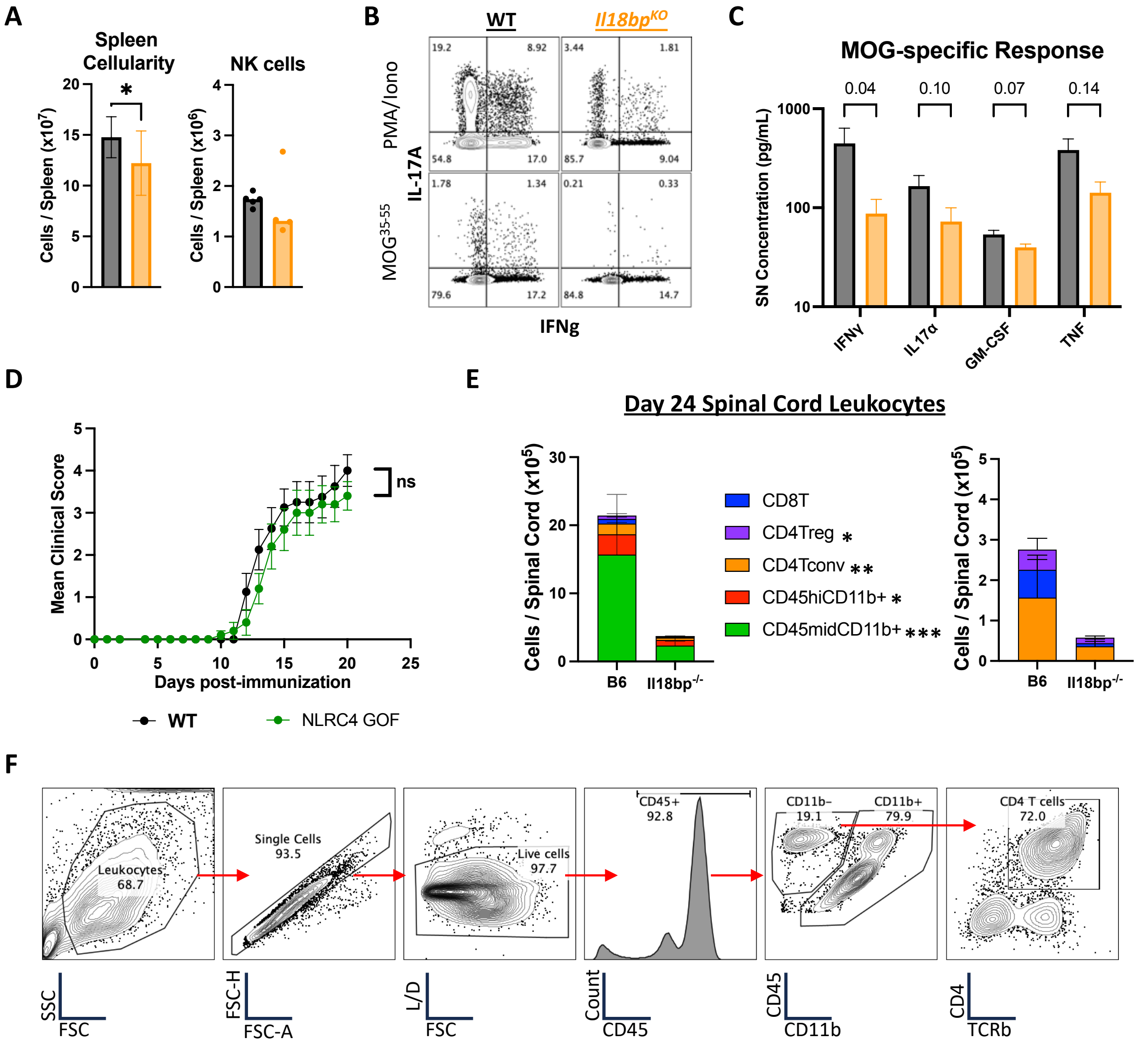


**Supplemental Figure 2: Excess IL-18 prevents accumulation of activated CD4Tauto in the periphery and CNS. (A)** Flow cytometric quantification of splenic leukocyte populations at day 12. **(B)** Representative flow plots of IFNg and IL-17A expression in day 12 splenic CD4Tconv following PMA/IONO and $MOG^{35\text{-}55}$ stimulation (quantified in Figure 2C). **(C)** Measurement of cytokine accumulation in culture supernatant of day 12 splenocytes stimulated with $MOG^{35\text{-}55}$. **(D)** *Nlrc4*$^{GOF}$ mice have an activating mutation in the NLRC4 inflammasome which leads to overproduction of IL-18 specifically by intestinal epithelial cells. Mean clinical score of WT (n=8) and *Nlrc4*$^{GOF}$ (n=10) mice is shown. **(E)** Flow cytometric quantification of spinal cord leukocytes on day 24. **(F)** Representative gating strategy of CD4 T-cells isolated from the spinal cord.

(A,B,E,F) Data representative of 2-3 independent experiments. (C,D) Data pooled from 2 experiments. Error bars = SEM. Statistical analysis: (A) unpaired t-test, (C,E) unpaired t-tests with Holm-Sidak correction of p-value, (D) Mann-Whitney of AUC. ns = not significant. *$p < 0.05$, **$P<0.01$, *** $P<0.001$.

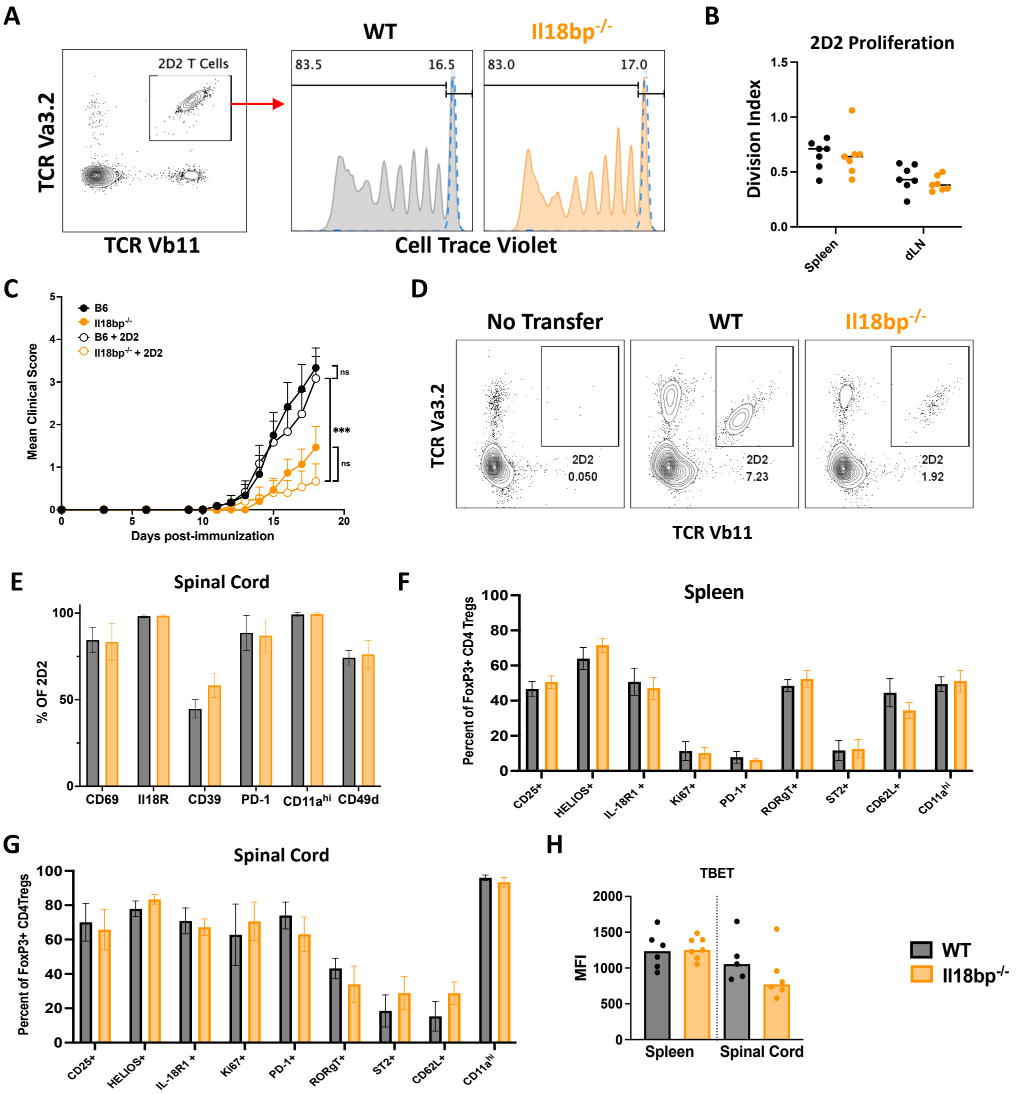


**Supplemental Figure 3: Excess IL-18 demonstrates defects in CD4T$_{auto}$ late in priming but no phenotypic differences in CD4T$_{reg}$. (A-B)** 4x10$^6$ CTV-labeled splenic CD4 T-cells from naïve 2D2 mice were transferred to WT or *Il18bp$^{KO}$* mice on day -1 followed by EAE induction on day 0. On day 5, 2D2 proliferation was assessed by CTV dye dilution via flow cytometry. **(A)** Representative 2D2 CD4 T-cell plots and CTV proliferation histograms plots. Dashed blue line represented no immunization control. **(B)** Calculated division index of CTV-labeled 2D2. **(C-E)** 2.5x10$^5$ splenic CD4 T-cells from naïve 2D2 mice were transferred to WT or *Il18bp$^{KO}$* mice on day -1 followed by EAE induction on day 0 (as in Figure 2F-I). **(C)** Mean clinical score of mice with 2D2 transfer (WT n=12, *Il18bp$^{KO}$* n=15) or without (WT n=12, *Il18bp$^{KO}$* n=15) and **(D)** detection of 2D2 T-cells in day 12 spleen is shown. **(E)** Surface protein expression on spinal cord 2D2 T-cells at day 18 (only shown for mice with CD4Ts above limit of detection). **(F,G)** Expression of surface markers and transcription factors on FOXP3$^+$CD4T$_{reg}$ from spleen and spinal cord of WT and *Il18bp$^{KO}$* mice at day 20. **(H)** MFI of Tbet, a transcription factor described in Th1 adaptation of CD4T$_{reg}$ cells.

(A-E) Data pooled from 2 experiments. (F-H) Data representative of 2 experiments. Error bars = SEM. Statistical analysis: (B,E-H) unpaired t-tests, p-value with Holm-Sidak correction (all comparisons not significant). (C) Kruskal-Wallis of AUC with Dunn's post-test of pairwise comparisons. ns = not significant, ***p<0.001.

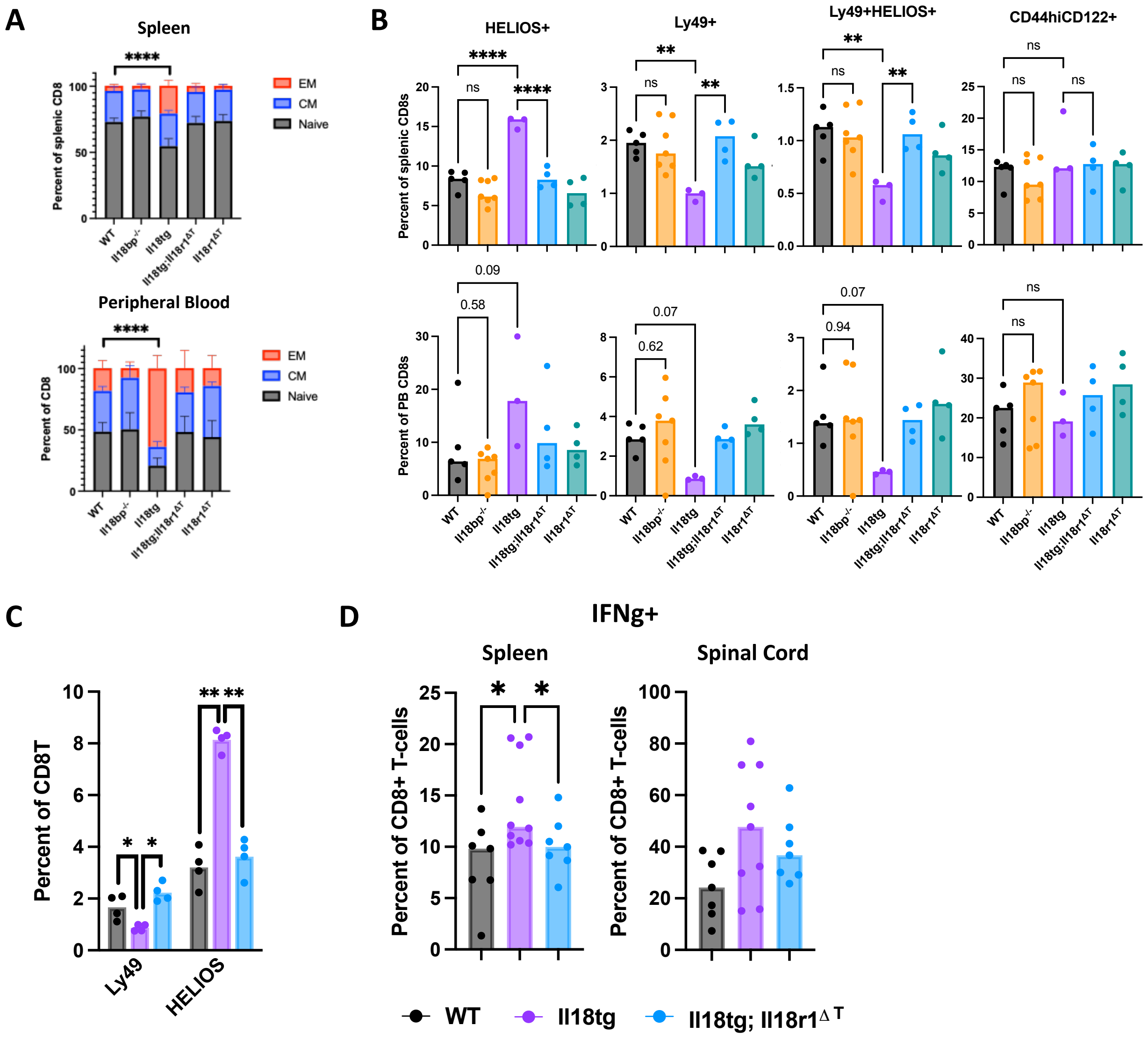


**Supplemental Figure 4: Chronic excess IL-18 decreases CD8 T-cell Ly49 expression but increases HELIOS expression and IFNg production.** Baseline flow cytometric staining of CD8 T-cells from spleen and peripheral blood in naïve mice of the indicated genotypes. **(A)** CD8 T-cells were differentiated into naïve ($CD44^{lo}CD62L^{+}$), central memory ($CD44^{hi}CD62L^{+}$), or effector memory ($CD44^{hi}CD62L^{-}$). **(B)** Total CD8 T-cells were interrogated for putative suppressive marker Ly49, HELIOS, and CD122. **(C)** Expression of Ly49 and HELIOS in total CD8 T-cells from dLN at day 20 in the indicated genotypes (described in Figure 3). **(D)** IFNg production by CD8 T-cells following 6h PMA stimulation and ICS at day 20.

(A-C) Representative data. (D) Pooled data from 2 experiments. Error bars = SEM. Statistical analysis: (A) One-way ANOVA of EM with Dunnett's post-test of pairwise comparisons to WT, (B-D) One-way ANOVA with Dunnett's post-test of pairwise comparisons to Il18tg (C, D) or the indicated groups (B). ns = not significant. *$p < 0.05$, **$p < 0.01$, ****$p < 0.0001$.

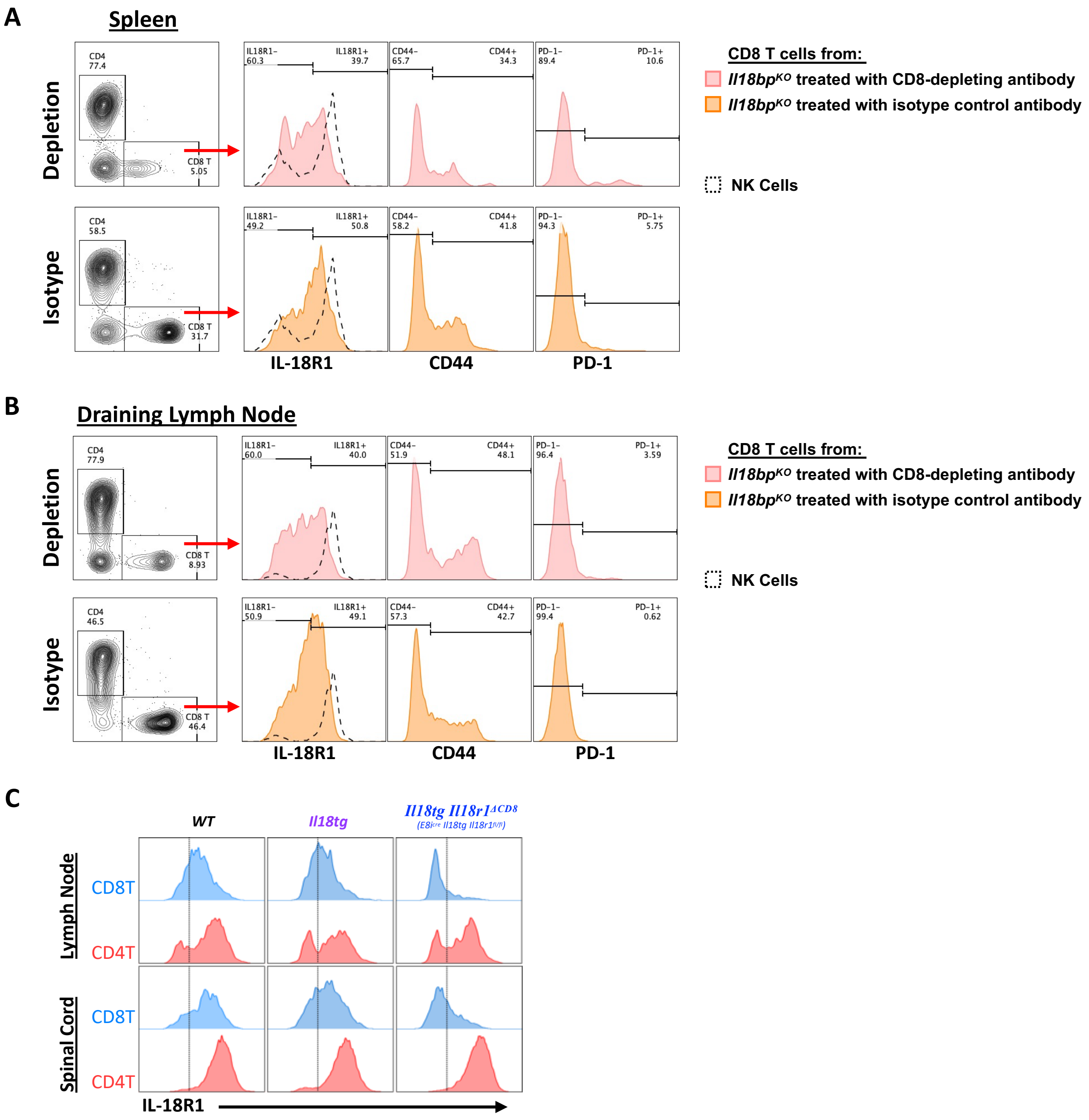


**Supplemental Figure 5: CD8 depletion or Tamoxifen treatment incompletely eliminates IL-18-receptor-expressing CD8 T-cells. (A,B)** Representative plots of CD4 and CD8 expression in total T-cells (repeated from Figure 5I) and histograms of IL-18R1, CD44, and PD-1 expression on CD8 T-cells from *Il18bp*$^{KO}$ mice receiving CD8 depleting or isotype control antibodies as in Figure 5H. Dashed line represents NK-cell IL-18R1 expression as a control. **(C)** Representative flow plots of IL-18R1 expression on CD44$^{hi}$ CD4 and CD8 T-cells from the indicated genotypes (as described in Figure 6) at day 23.

(A-C) Data are representative of 2-3 experiments.

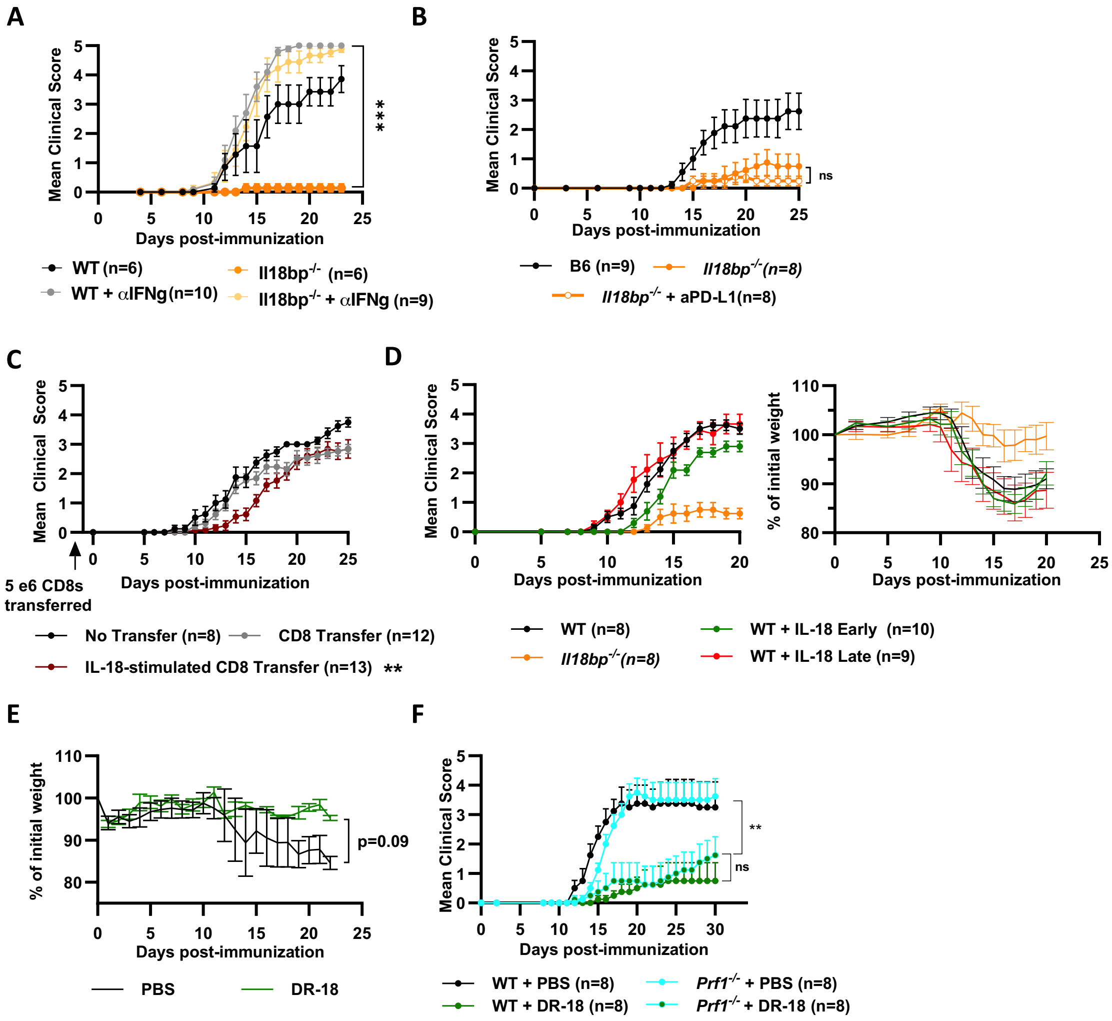


**Supplemental Figure 6: Excess IL-18 requires IFNg but not PD-L1 or perforin for protection.** Mean EAE clinical score of WT or *Il18bp*$^{KO}$ mice treated with **(A)** an IFNg-neutralizing antibody (200ug, days 0, 4, and 8); or **(B)** an PD-L1 blocking antibody (250ug, every 3 days) beginning day 0. **(C)** Splenic CD8 T-cells were isolated from day 10 WT mice immunized against MOG$^{35\text{-}55}$ and cultured for 3 days with or without IL-18 (50 pg/mL). Mock and IL-18 treated CD8 T-cells were transferred to naïve WT mice on day -1 followed by EAE induction on day 0. Mean EAE clinical score over time is shown. **(D)** Mean EAE clinical score and weight change of *Il18bp*$^{KO}$ and WT mice treated with recombinant IL-18 (1ug, every other day) starting at day 1 (early) or day 9 (late). Control mice received PBS treatment throughout. **(E)** Weight change following EAE induction of WT mice receiving PBS or DR-18 (2ug every 3 days, from days 0 to 15, as described in Figure 7A). **(F)** Mean EAE clinical score of WT and *Prf1*$^{KO}$ mice treated with PBS or DR-18 (2ug ever 3 days, from day 0 to 30).

(A-D, F) Data pooled from 2 experiments. Number of mice (n) indicated in individual legends. (E) Data representative of 2 experiments. Error bars = SEM. Statistical analysis: (A-D,F) Kruskal-Wallis of AUC with Dunn's post-test of pairwise comparisons within genotypes (A, B, D), all groups (C), or shown comparisons only (F). (E) 2-way RM ANOVA, p-value of treatment effect. ns=not significant, **p<0.01, ***p<0.001, ****p<0.0001.

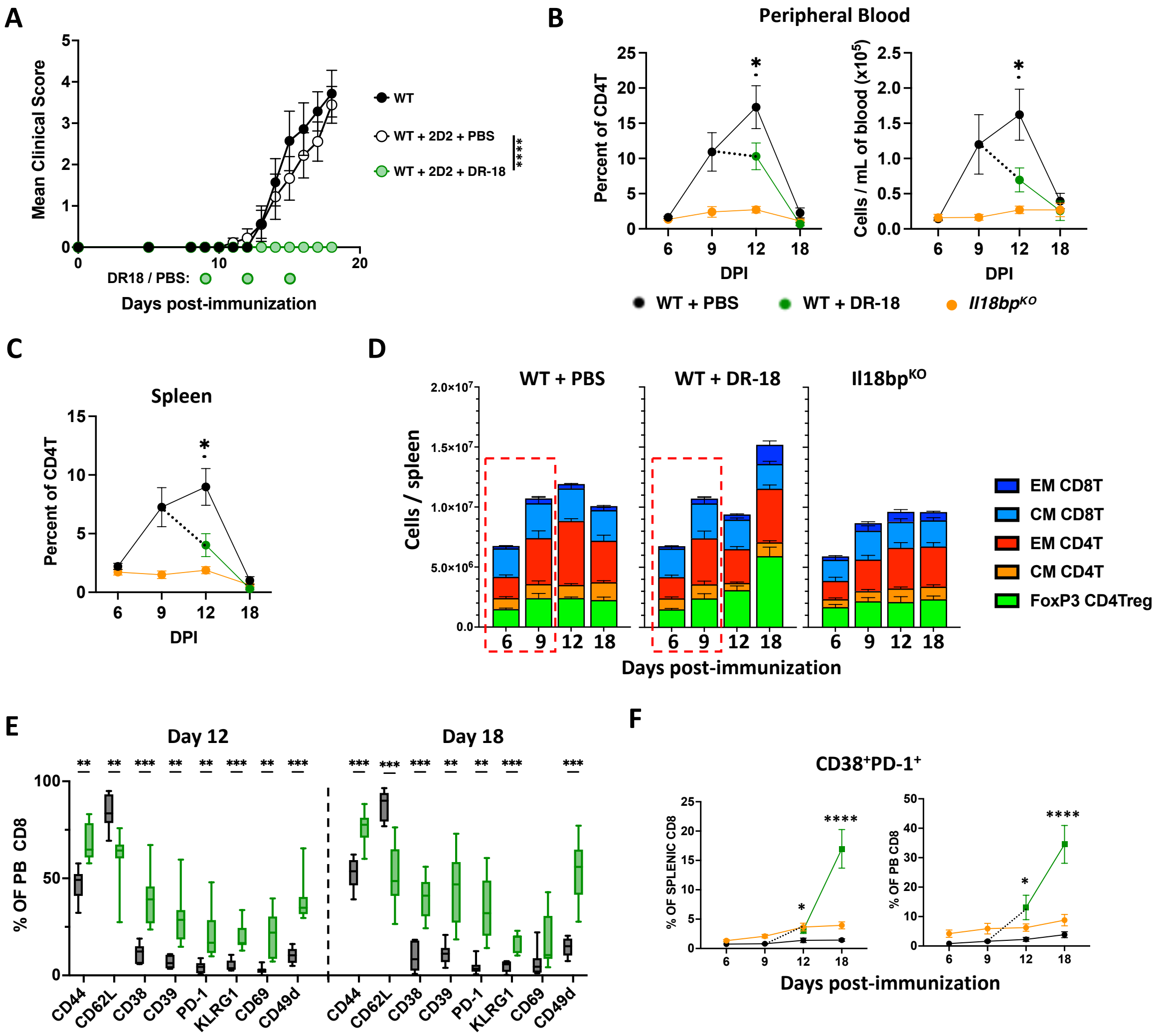


**Supplemental Figure 7: DR-18 disrupts autoimmune effector activity and protects from EAE immunopathology.** Extended data from Figure 7. 2.5 x$10^5$ splenic CD4 T-cells from naïve 2D2 mice were transferred to WT and *Il18bp*$^{KO}$ mice on day -1. Following EAE induction on day 0, half of the WT mice were randomized to receive DR-18 or PBS on days 9, 12, & 15. **(A)** Mean clinical score of WT mice either untreated (no transfer) or receiving 2D2 T-cell transfer, with PBS or DR-18 treatment (n=8 for all groups). **(B,C)** Quantification of 2D2 T-cells in peripheral blood and spleen at various time points. **(D)** Quantification of splenic $CD4T_{conv}$ and CD8 T-cell subsets (central memory, $CD44^+CD62L^+$; effector, $CD44^+CD62L^-$) along with CD4Tregs ($FOXP3^+$) over time. Data in red box represent replicate data from day 6 and 9 WT mice prior to treatment. **(E)** Expression of surface markers on peripheral blood CD8 T-cells from WT mice treated with PBS or DR-18. **(F)** Percent of dual-expressing $CD38^+PD\text{-}1^+$ CD8 T-cells in spleen and PB over time.

Data pooled from 2 experiments. Error bars = SEM except for box and whisker plot. Statistical analysis: (A) Kruskal-Wallis of AUC with Dunn's post-test of pairwise comparisons (B, C, F) unpaired t-tests of WT vs WT+DR-18 on days 12 and 18, p-value with Holm-Sidak correction, (D) statistical analysis performed in main figure, complete data provided for context, (E) unpaired t-tests with Holm-Sidak correction of p-value. Only $p_{adj}<0.05$ is shown. *$p< 0.05$, **$p<0.01$, ***$p<0.001$, ****$p<0.0001$.
